# Microglial lipid signaling drives glioblastoma invasion and represents a therapeutic vulnerability

**DOI:** 10.64898/2026.04.24.720633

**Authors:** Carla Pallarés-Moratalla, Mélanie Guyot, Krish Skandha Gopalan, Bernhard Robl, Jakub Idkowiak, Jonas Dehairs, Nina Ravoet, Kinga Anna Nowak, Nikolina Dubroja Lakić, Linqian Weng, Vasileios Konstantakos, Xander Spotbeen, Sarah-Maria Fendt, Stein Aerts, Frederik De Smet, Johannes V. Swinnen, Gabriele Bergers

**Affiliations:** Laboratory of Tumor Microenvironment and Therapeutic Resistance, VIB-KU Leuven Center for Cancer Biology, Leuven, Belgium; Department of Oncology, KU Leuven, Leuven, Belgium; Department of Pediatrics and Adolescent Medicine, Comprehensive Cancer Center and Comprehensive Center for Pediatrics, Medical University of Vienna, Vienna, Austria; Laboratory of Lipid Metabolism and Cancer, KU Leuven, Leuven, Belgium; Laboratory for Precision Cancer Medicine, Translational Cell and Tissue Research Unit, Department of Imaging & Pathology, KU Leuven, Leuven, Belgium; KU Leuven Institute for Single Cell Omics (LISCO), Leuven, Belgium; Laboratory of Computational Biology, VIB Center for AI & Computational Biology (VIB.AI), Leuven, Belgium; VIB-KU Leuven Center for Brain & Disease Research, Leuven, Belgium; Department of Human Genetics, KU Leuven, Leuven, Belgium; Laboratory of Cellular Metabolism and Metabolic Regulation, VIB Center for Cancer Biology, VIB, Leuven, Belgium; Leuven Cancer Institute (LKI), Leuven, Belgium

## Abstract

Glioblastoma (GBM) is characterized by diffuse infiltration into the surrounding brain, which precludes complete surgical resection, the strongest determinant of patient survival. The mechanisms that drive this invasive growth remain incompletely understood. Here we identify a lipid-mediated paracrine signaling axis through which microglia, the resident macrophages of the brain, promote glioma invasion. Integrating single-cell transcriptomics, spatial lipidomics, and functional perturbation across mouse models and human GBM samples, we show that invading tumor cells engage and reprogram microglia via CSF1R–PI3K signaling. This induces a metabolic switch in microglia, leading to the secretion of bioactive lipids, including lysophosphatidylcholines (LPCs) and lysophosphatidic acids (LPAs), which act as pro-invasive cues across GBM subtypes through distinct downstream pathways. Disruption of the microglia–GBM axis, either by inhibiting CSF1R signaling or by blocking lipid mobilization, reduces lipid secretion and suppresses tumor invasion. Targeting downstream LPA–LPAR or YAP/TAZ signaling further constrains invasion in a context-dependent manner. Together, these findings define a lipid-driven signaling circuit that links the tumor microenvironment to glioma invasion and identify therapeutic strategies to limit tumor infiltration and improve surgical resectability.

## Introduction

Glioblastoma (GBM) is the most aggressive primary brain tumor and remains uniformly fatal despite multimodal therapy. Its diffuse infiltration into the surrounding brain parenchyma precludes complete surgical resection, the strongest predictor of survival, resulting in a median survival of approximately 15 months^1–3^. Targeting the mechanisms that drive tumor invasion therefore represents a critical unmet need to improve resectability and patient outcomes^4^.

GBM cells infiltrate the brain by commonly co-opting existing anatomical structures, including white matter tracts, meninges and ventricular linings, extending along myelinated fiber tracts and basement membranes of blood vessels to distant sites in the brain parenchyma or the leptomeningeal space^5^. During invasion, glioma cells engage in dynamic interactions with various resident host cells in the tumor microenvironment and surroundings, receiving signals that promote migration, survival, and resistance to therapy^2,4,6^. Among these are microglial cells (MGs), abundant brain-resident macrophages that serve as immune sentinels, participating in both innate and adaptive immune responses and are distributed throughout the entire brain^7^. Peripheral macrophages together with microglia can comprise up to 30-50% of tumor-associated macrophages (TAMs) in GBMs^8^. Microglia contribute to GBM progression by promoting angiogenesis, invasion, immunosuppression, and tumor growth^4,6^. Colony-stimulating factor-1 receptor (CSF1R) signaling is a central regulator of microglial/macrophage activation and has been implicated in shaping a tumor-permissive microenvironment, in part through PI3K-dependent programs that support tissue remodeling and immunosuppression^9^. Microglia and macrophages can promote tumor progression through the secretion of cytokines, growth factors, and direct cell–cell interactions^10–12^. The contribution of lipids, however, has remained largely unexplored.

Lipids are essential regulators of cellular homeostasis and tumor biology, serving as structural components, energy sources, and signaling molecules. In multiple cancers, lipid metabolism supports tumor growth, epithelial–mesenchymal transition, and metastatic dissemination^13–18^. Within the brain, lipids constitute a major fraction of tissue mass and play critical roles in cellular communication and function^18,19^. Cancer cells can acquire lipids through *de novo* synthesis or from the microenvironment ^20–22^, however, the role of microenvironment-derived lipids in GBM invasion is less explored.

Here, we identify a lipid-mediated paracrine signaling axis through which microglia drive glioma invasion. Integrating spatial lipidomics and proteomics with single-cell RNA sequencing and functional perturbation, we show that GBM-associated microglia secrete bioactive lipids, including lysophosphatidylcholines (LPCs) and lysophosphatidic acids (LPAs), that directly promote tumor cell migration independently of genetic background. Mechanistically, CSF1R–PI3K signaling regulates lipid mobilization in microglia, and its inhibition reprograms microglia toward a tumor-inhibiting state characterized by intracellular lipid accumulation and reduced lipid secretion, thereby limiting tumor invasion across multiple models.

## Results

### Reciprocal activation of GBM and microglia promote tumor invasion

To define functional interactions between infiltrating GBM cells and MGs, we employed two syngeneic murine GBM (EGFRvIII^+^ NSCG and NF1/p53/Pten-deficient NFpp10) and four human GBM xenografts of which two display EGFR amplifications or mutations (LBT012 and LBT090). Every GBM exhibited a close spatial association with microglial cells at the invasive front (**Figure 1A; Figures S1A and S1B**). The glioma cells further displayed a widespread infiltration of single or clustered MG-associated GBM cells throughout the entire brain (**Figure 1A**; **Figures S1A and S1B**).

**Figure 1.**
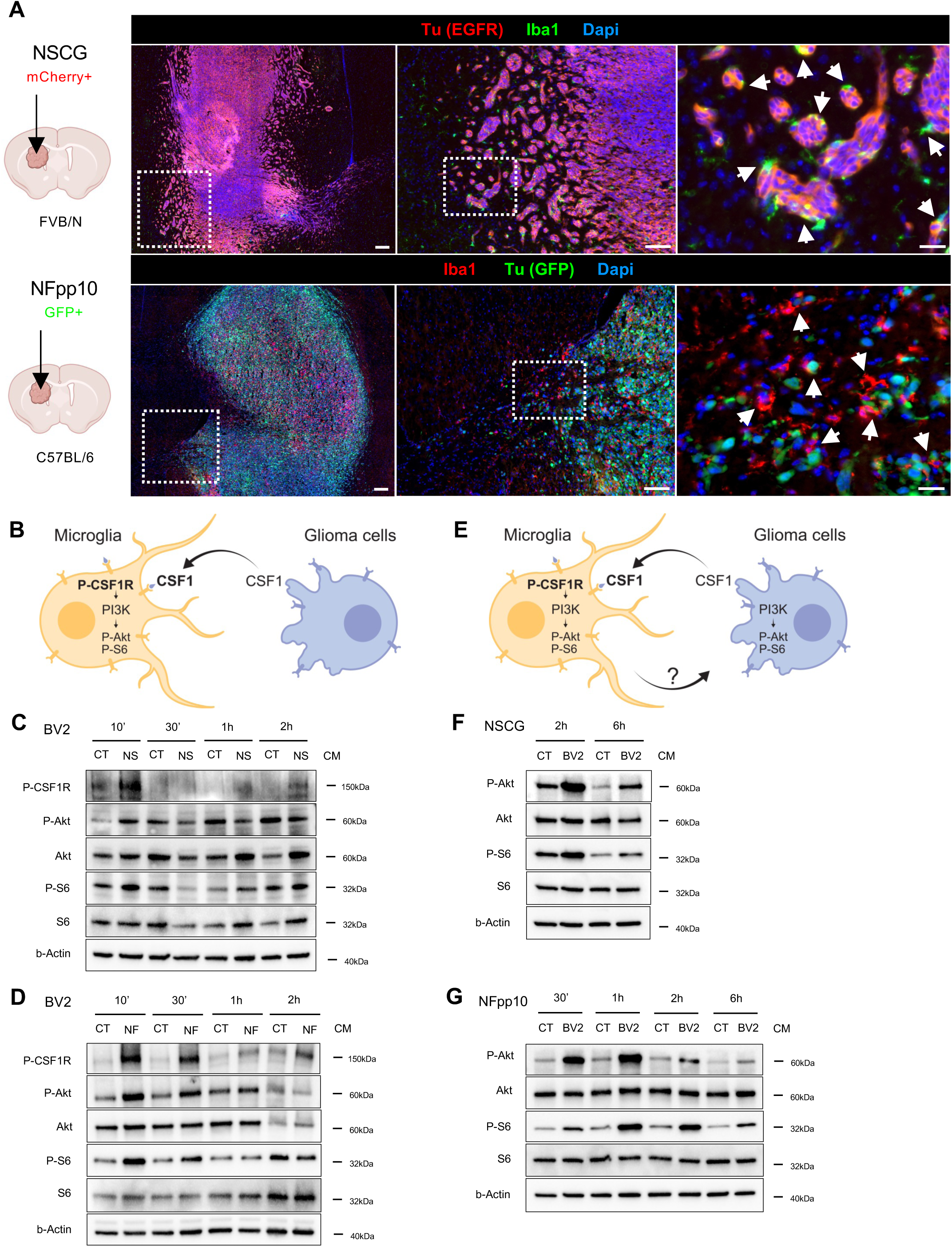
Spatial association and activation of MGs and GBM cells at the tumor rim. (A) Immunofluorescence images of NSCG (upper panel, EGFR^+^, red) and NFpp10 (lower panel, GFP^+^, green) tumor cells associated with microglial cells (Iba1^+^, green up, red down) at the invasive rim of tumor-bearing mice. Scales (from left to right) 200µm, 100µm, and 20µm. (B) Schematic of the MG-GBM crosstalk Part I. (C) Representative immunoblots of P-CSF1R, P-Akt, and P-S6 levels in murine BV2 microglial cells stimulated for 10 minutes, 30 minutes, 1 hour, or 2 hours with NSCG conditioned-media. (D) Representative immunoblots of P-CSF1R, P-Akt, and P-S6 levels in murine BV2 microglial cells stimulated for 10 minutes, 30 minutes, 1 hour, or 2 hours with NFpp10 conditioned-media. (E) Schematic of the MG-GBM crosstalk Part II. (F) Representative immunoblots of P-Akt, and P-S6 levels in NSCG cells stimulated for 2 hours or 6 hours with BV2 conditioned-media. (G) Representative immunoblots of P-Akt, and P-S6 levels in NFpp10 cells stimulated for 30 minutes, 1 hour, 2 hours or 6 hours with BV2 conditioned-media.

Coculture experiments of murine BV2 microglial cells treated with conditioned media (CM) of NSCG or NFpp10 GBM cells displayed increased CSF1R (P-CSF1R) and downstream PI3K activation (P-Akt, P-S6) (**Figures 1B-1D**). Interestingly, BV2-conditioned NSCG and NFpp10 cells also displayed increased Akt and S6 kinase activation (**Figures 1E-1G**), suggestive of a reciprocal crosstalk between GBM and microglial cells via mutual PI3K/Akt/mTOR pathway activation, known to promote cell migration and survival cues.

Congruent with these results, we found that murine BV2 MG cells substantially increased the invasive capacity of both NSCG and NFpp10 cells *in vitro* (**Figure 2A**). Likewise, human HMC3 microglia enhanced invasion across two human EGFR amplified/mut^+^ patient-derived glioma lines (LBT012 and LBT090) and four EGFR wildtype cell lines (LBT065, LBT007, LBT059, and LBT123) (**Figures 2B and 2C, Figure S2A**). Moreover, consistent with the GBM-induced CSF1R/PI3K activation in microglia, pharmacological inhibition of CSF1R with PLX3397 (PLX) or PI3Kγ/δ with IPI-145 (IPI) in BV2 and HMC3 microglial cells, significantly reduced invasion of murine and human GBMs *in vitro* (**Figures 2D-2F, Figure S2B**).

**Figure 2.**
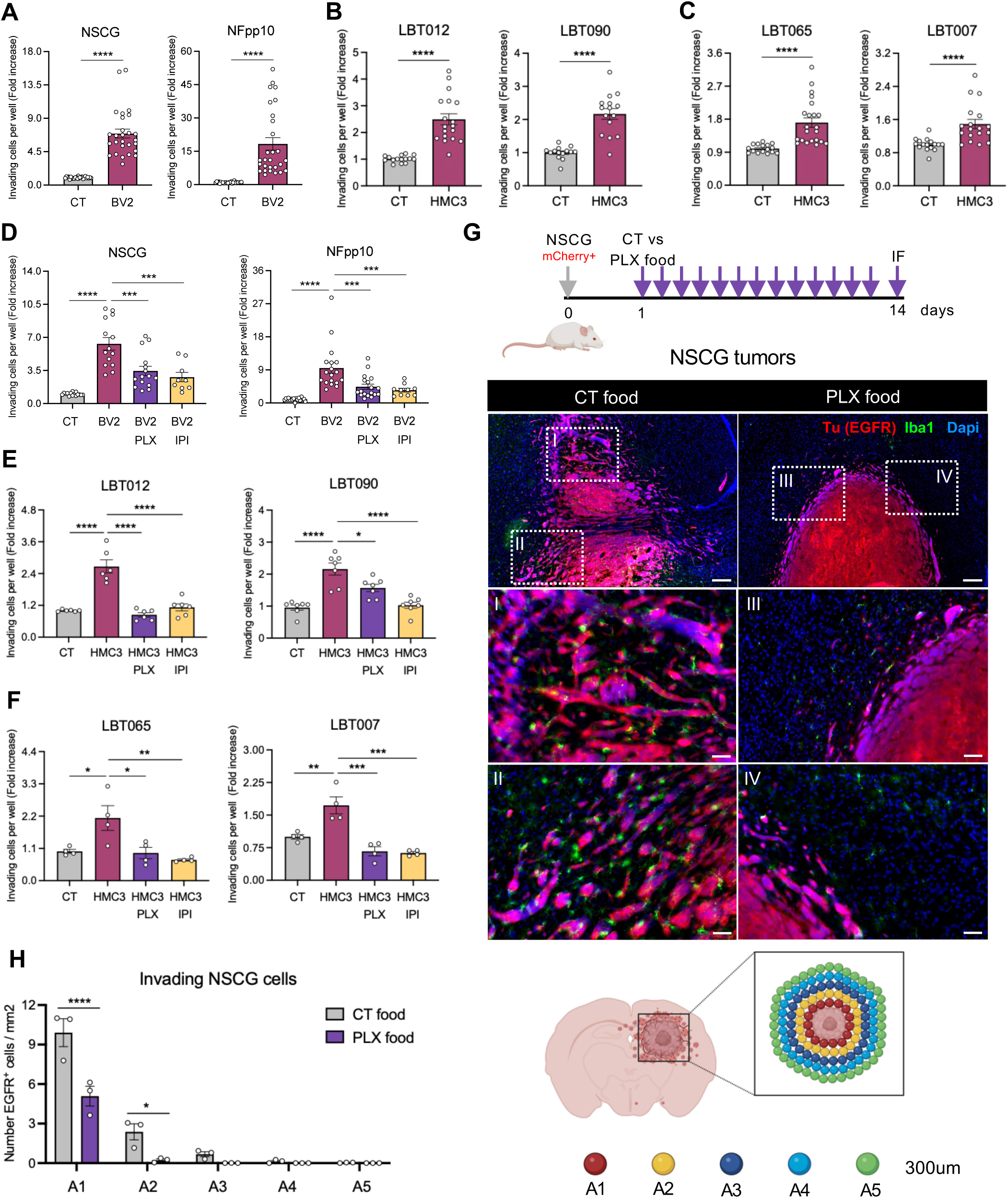
GBM invasion relies on the microglial cell activation state. (A) Invasion assays of NSCG (left) and NFpp10 (right) cells in the presence or absence of murine BV2 microglial cells. Fold increase. NSCG: n=10, Mann-Whitney test (T-test non-parametric), **** p-value<0.0001; NFpp10: n=10, Mann-Whitney test (T-test non-parametric), **** p-value<0.0001. (B) Invasion assays of LBT012 LamB (left) and LBT090 Lami (right) cells (EGFR amplified/mut^+^) in the presence or absence of human HMC3 microglial cells. Fold increase. LBT012 LamB: n=12, Mann-Whitney test (T-test non-parametric), **** p-value<0.0001; LBT090 Lami: n=6, Mann-Whitney test (T-test non-parametric), **** p-value<0.0001. (C) Invasion assays of LBT065 LamB (left) and LBT007 Lami (right) cells (EGFR wildtype) in the presence or absence of human HMC3 microglial cells. Fold increase. LBT065 LamB: n=11, Mann-Whitney test (T-test non-parametric), **** p-value<0.0001; LBT007 Lami: n=9, Mann-Whitney test (T-test non-parametric), **** p-value<0.0001. (D) Invasion assays of NSCG (left) and NFpp10 (right) cells in the presence or absence of murine BV2 microglial cells, PLX3397, and/or IPI-145. Fold increase. NSCG: n=5, One-way ANOVA, **** p-value<0.0001, *** p-value<0.003; NFpp10: n=7, One-way ANOVA, **** p-value<0.0001, *** p-value<0.007. (E) Invasion assays of LBT012 LamB (left) and LBT090 Lami (right) cells (EGFR amplified/mut^+^) in the presence or absence of human HMC3 microglial cells, PLX3397, and/or IPI-145. Fold increase. LBT012 LamB: n=6, One-way ANOVA, **** p-value<0.0001; LBT090 Lami: n=4, One-way ANOVA, * p-value=0.0147, **** p-value<0.0001. (F) Invasion assays of LBT065 LamB (left) and LBT007 Lami (right) cells (EGFR wildtype) in the presence or absence of human HMC3 microglial cells, PLX3397, and/or IPI-145. Fold increase. LBT065 LamB: n=4, One-way ANOVA, * p-value<0.0226, ** p-value=0.0050; LBT007 Lami: n=4, One-way ANOVA, ** p-value=0.0038, *** p-value<0.0002. (G) Representative immunofluorescent images of invading NSCG cells (EGFR^+^, red) and their association with microglial cells (Iba1^+^, green) in control- (left panel) and PLX5622-treated (right panel) tumor bearing mice. Scales (upper panels to medium/lower panels): 200µm, 50µm. (H) Quantification and representative scheme of invading NSCG cells per areas of 300µm from the tumor rim in control- and PLX5622-treated tumor bearing mice. n=3, 2way ANOVA, **** p-value<0.0001. All data represent mean ± SEM.

To validate these findings *in vivo*, NSCG cells were injected intracranially into FVB/N mice, followed by administration of PLX3397-formulated or control chow for 13 days, a duration sufficient to induce microglia reprogramming and tumor invasion. Consistent with our *in vitro* results, PLX treatment significantly suppressed glioma invasion (**Figures 2G and 2H**) and reduced microglial abundance in the brains of tumor-bearing mice (**Figure S2C**).

Together, these findings reveal that GBM cells activate CSF1R and downstream PI3K pathways in microglia which in turn secrete molecules that promote GBM invasion. Pharmacological blockade of microglia activity is sufficient to impair the invasive capacity of both murine and human GBM cells.

### GBM-associated microglia undergo lipid metabolic reprogramming

What are then the MG-derived mediators that drive GBM invasion? We performed single-cell RNA sequencing (scRNA-seq) of isolated glioma and myeloid cells from the tumor rim and core of naïve and PLX-treated NSCG and NFpp10-bearing mice to identify pathways in tumor and microglial cells that were prevalent at the invasive front (**Figure 3A, Figures S3A and S3B**).

**Figure 3.**
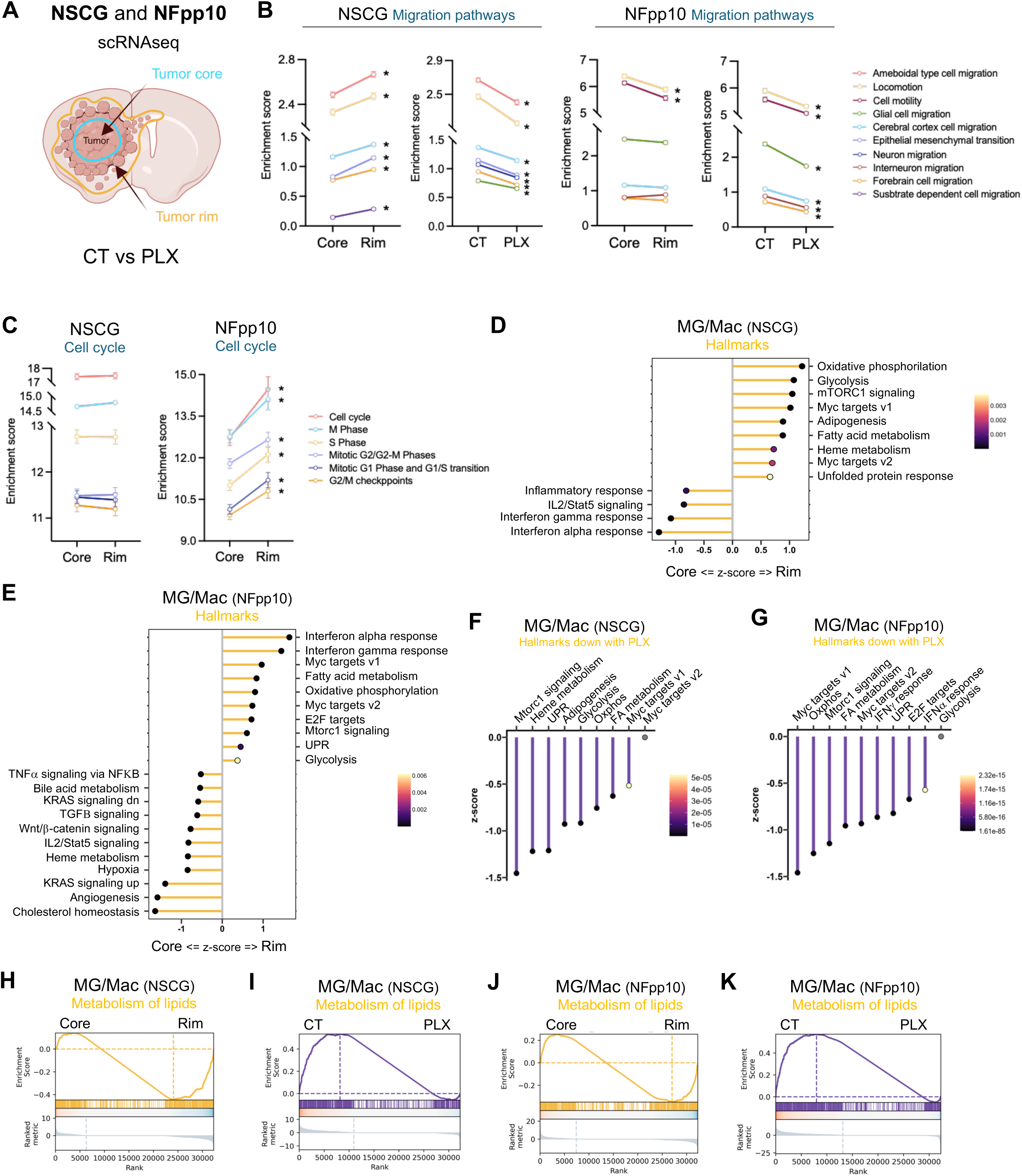
GBM-associated MG undergo lipid metabolism reprogramming. (A) Schematic of the NSCG and NFpp10 scRNAseq samples from untreated and PLX-treated tumor bearing mice. (B) Line plot of migration-related pathways in NSCG and NFpp10 tumor core-rim, and tumor rim control-PLX treated samples. Mean ± SEM. * p-value<0.0001. (C) Line plot of cell cycle-related pathways in NSCG and NFpp10 tumor core-rim untreated samples. Mean ± 95% CI. * p-value<0.0001. (D) Lollipop chart of the most significant hallmarks in the microglia/macrophages of NSCG-tumor core and rim samples. (E) Lollipop chart of the most significant hallmarks in the microglia/macrophages of NFpp10-tumor core and rim samples. (F) Lollipop chart of the most significant hallmarks in the microglia/macrophages of NSCG-tumor rim brains upon PLX-treatment. (G) Lollipop chart of the most significant hallmarks in the microglia/macrophages of NFpp10-tumor rim brains upon PLX-treatment. (H) Leading edge plot of the metabolism of lipids reactome in microglia/macrophages of NSCG tumor core and rim samples. p-adj=2.4E-07. NES=2.285288. (I) Leading edge plot of the metabolism of lipids reactome in microglia/macrophages of NSCG tumor rim control and PLX-treated samples. p-adj=3.57E-07. NES=0.7191733. (J) Leading edge plot of the metabolism of lipids reactome in microglia/macrophages of NFpp10 tumor core and rim samples. p-adj=2.3E-02. NES=0.467985. (K) Leading edge plot of the metabolism of lipids reactome in microglia/macrophages of NFpp10 tumor rim control and PLX-treated samples. p-adj=8.17E-45. NES=1.686654.

As expected, transcriptomic analysis revealed enrichment of migration-associated pathways in NSCG tumor rim samples relative to the tumor core (**Figure 3B**), including pathways linked to amoeboid cell migration, locomotion, cerebral cortex cell migration, and glial cell migration (**Figure 3B**). These invasion-related programs were broadly suppressed in NSCG rim samples following PLX treatment (**Figure 3B**).

Interestingly, NFpp10 tumor rim cells did not exhibit a significant upregulation of the majority of migration-associated pathways observed in NSCG rims (**Figure 3B**). PLX treatment, however, also downregulated distinct migration-related pathways in NFpp10 GBM, consistent with the response observed in NSCG tumors (**Figure 3B**). Notably, the NFpp10 tumor cells at the invasive rim exhibited a pronounced increase of cell-cycle associated pathways (**Figure 3C**), suggestive of a proliferation-dominant phenotype in NFpp10 cells. This proliferative bias was not observed in NSCG tumors (**Figure 3C**).

Next, we performed a detailed transcriptomic analysis of microglia and macrophage populations (MG/Mac) across tumor core and rim regions from NSCG and NFpp10 GBMs (**Figures 3D and 3E**). NSCG rim-located MG/Mac exhibited robust activation of specific metabolic programs, including oxidative phosphorylation, glycolysis, and lipid-associated pathways such as adipogenesis and fatty acid metabolism, whereas core-associated MG/Mac were enriched for inflammatory response signatures, including interferon-γ and interferon-α signaling (**Figure 3D**). A comparable metabolic phenotype was observed in MG/Mac from NFpp10 tumor rims, marked by upregulation of fatty acid metabolism, oxidative phosphorylation, and glycolysis (**Figure 3E**). In contrast, MG/Mac from NFpp10 tumor cores preferentially activated inflammatory pathways, including TNFα, TGFβ, and IL2/STAT5 signaling, while interferon-α and -γ responses were enriched at the tumor rim (**Figure 3E**).

These findings suggest that tumor rim-associated MG/Mac undergo a pronounced metabolic reprogramming and polarize toward a lipid-metabolically active state. In line with this interpretation, PLX treatment markedly suppressed these metabolic programs in MG/Mac from both NSCG and NFpp10 tumor rims (**Figures 3F and 3G**) and concomitantly, decreased PI3K/AKT/mTOR signaling, a pathway linked to immunosuppressive MG/Mac states (**Figures S3C and S3D**). Reactome pathway analysis further corroborated these findings, revealing selective upregulation of lipid metabolism pathways in MG/Mac from NSCG and NFpp10 tumor rims (**Figures 3H and 3J**), which was abrogated upon PLX treatment (**Figures 3I and 3K**). Given this consistent metabolic shift, we next examined lipid-associated functional programs in greater detail. MG/Mac populations from NSCG and NFpp10 tumor rims displayed enrichment of pathways involved in lipid remodeling and trafficking, including lipid export, localization, storage, and cellular response to lipids (**Figures S3E and S3F**). In contrast, PLX treatment downregulated lipid biosynthetic and metabolic processes in rim-associated MG/Macs across both GBMs (**Figures S3E and S3F**).

Collectively, our findings show that NSCG- or NFpp10-associated microglia underwent metabolic reprogramming marked by activated lipid metabolism, export, and lipid-responsive signaling pathways. CSF1R inhibition suppressed these lipid-associated programs, leading to reduced tumor invasion. These data suggest microglia-derived lipid synthesis, trafficking, and export as paracrine drivers of GBM invasion.

### GBM-activated microglia cells drive glioblastoma invasion via lipid secretion

Indeed, lipids secreted by MG proved essential for glioma invasion because depleting lipids from BV2-conditioned media (MG-CM) using a lipid removal agent (LRA) abolished its proinvasive effect. Unlike BV2 MG-CM, lipid-depleted BV2 MG-CM failed to promote invasion of both NSCG and NFpp10 tumor cells (**Figure 4A**). Importantly, this loss of promoting invasiveness was consistent across GBM models, including LRA-treated HMC3 MG-CM in human EGFR amplified/mut^+^ GBM cells (LBT012 and LBT090) (**Figure 4B**) and EGFR wildtype GBM cell lines (LBT065, LBT007, LBT059, and LBT123) (**Figure 4C, Figure S4A**).

**Figure 4.**
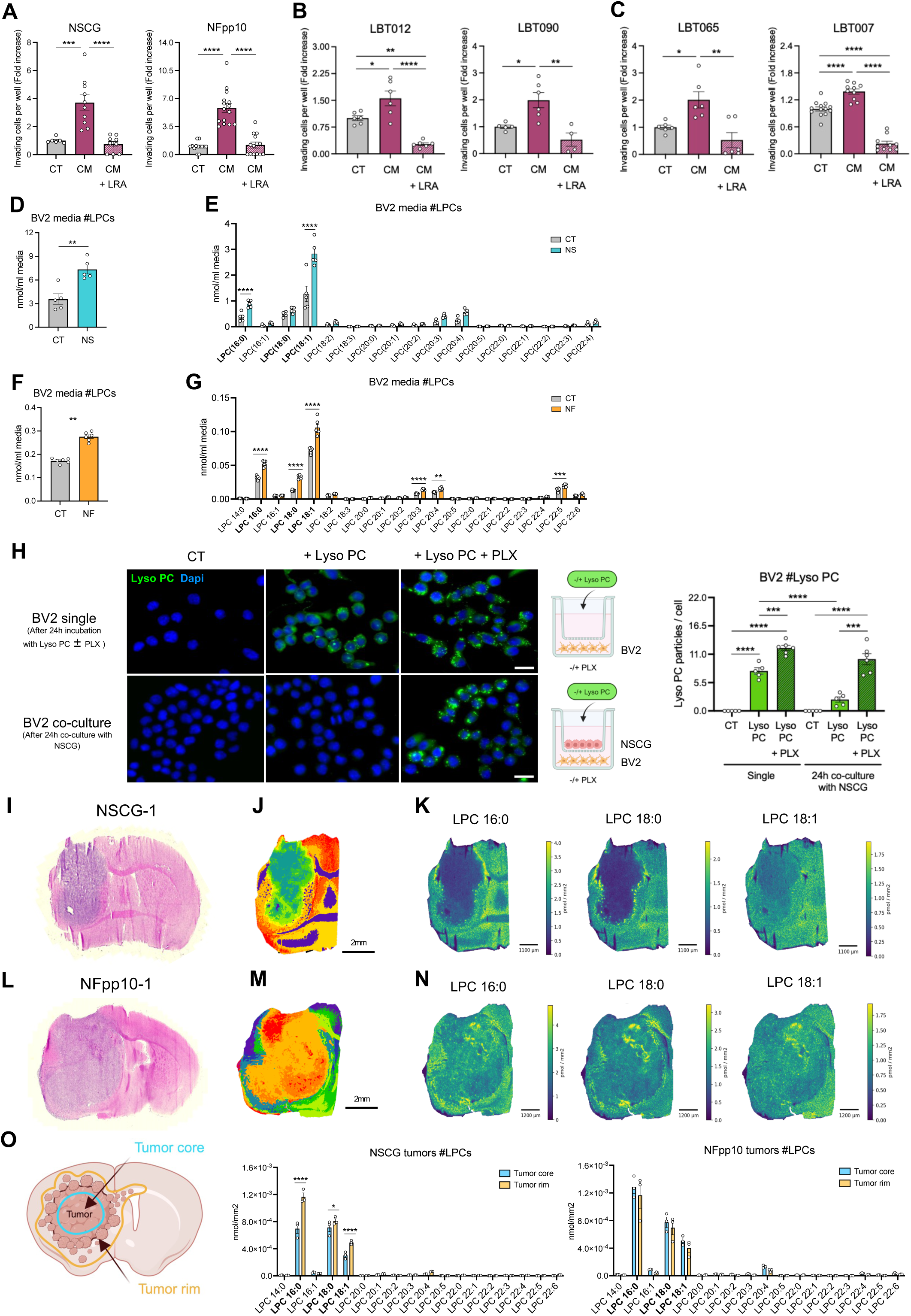
Microglia-secreted lipids drive GBM invasion. (A) Invasion assays of NSCG (left) and NFpp10 (right) cells in the presence or absence of BV2 conditioned-media (CM) with or without lipid removal agent (LRA). Fold increase. NSCG: n=4, One-way ANOVA, *** p-value=0.0004, **** p-value<0.0001; NFpp10: n=5, One-way ANOVA, **** p-value<0.0001. (B) Invasion assays of LBT012 LamB (left) and LBT090 Lami (right) cells (EGFR amplified/mut^+^) in the presence or absence of human HMC3 conditioned-media (CM) with or without lipid removal agent (LRA). Fold increase. LBT012 LamB: n=3, One-way ANOVA, * p-value=0.0194, ** p-value=0.0029, **** p-value<0.0001; LBT090 Lami: n=3, One-way ANOVA, * p-value=0.0203, ** p-value=0.0022. (C) Invasion assays of LBT065 LamB (left) and LBT007 Lami (right) cells (EGFR wildtype) in the presence or absence of human HMC3 conditioned-media (CM) with or without lipid removal agent (LRA). Fold increase. LBT065 LamB: n=3, One-way ANOVA, * p-value=0.0239; **p-value=0.0015; LBT007 Lami: n=4, One-way ANOVA, ** p-value=0.0068, **** p-value<0.0001. (D) Quantification of the sum of LPC species in BV2 media in the absence or presence of NSCG co-culture. n=5. One-way ANOVA, ** p-value=0.0079. (E) Quantification of the distinct LPC species in BV2 media in the absence or presence of NSCG co-culture. n=5. Two-way ANOVA, **** p-value<0.0001. (F) Quantification of the sum of LPC species in BV2 media in the absence or presence of NFpp10 co-culture. n=6. One-way ANOVA, ** p-value=0.0022. (G) Quantification of the distinct LPC species in BV2 media in the absence or presence of NFpp10 co-culture. n=6. Two-way ANOVA, ** p-value=0.0074, *** p-value=0.0005, **** p-value<0.0001. (H) Representative images and quantification of BV2 cells single or co-cultured with NSCG cells during 24 hours in the absence or presence of TopFluor Lyso PC, and/or PLX3397. Scale bar 30µm. n=3. One-way ANOVA, *** p-value=0.0003, **** p-value<0.0001. (I) Hematoxylin & Eosin (H&E) staining of an NSCG tumor brain (NSCG-1). (J) K-means lipid clustering of LPC, PC, PC P-/PC O-, SM, CAR in a NSCG tumor brain sample (NSCG-1) showing different lipid profiles in distinct areas. Signals normalized to their internal standard. Each color represent one lipid cluster. Scale 2mm. (K) Quantitative Mass Spectrometry Imaging (qMSI) of LPC 16:0, LPC 18:0, and LPC 18:1 in an NSCG tumor brain (NSCG-1). Scale 1100µm. (L) Hematoxylin & Eosin (H&E) staining of an NFpp10 tumor brain (NFpp10-1). (M) K-means lipid clustering of LPC, PC, PC P-/PC O-, SM, CAR in a NFpp10 tumor brain sample (NFpp10-1) showing different lipid profiles in distinct areas. Signals normalized to their internal standard. Each color represent one lipid cluster. Scale 2mm. (N) Quantitative Mass Spectrometry Imaging (qMSI) of LPC 16:0, LPC 18:0, and LPC 18:1 in an NFpp10 tumor brain (NFpp10-1). Scale 1200µm. (O) Quantification of the distinct LPC species in NSCG (left) and NFpp10 (right) tumor core and tumor rim tissue samples dissected using laser-capture microdissection followed by bulk lipidomics. NSCG: n=3, 2way ANOVA, * p-value=0.0229, **** p-value<0.0001; NFpp10, n=3, Two-way ANOVA, non-significant. All data represent mean ± SEM.

To identify the specific lipids driving GBM invasion, we performed bulk lipidomic profiling of BV2 MG-cells and their conditioned media (MG-CM) under mono- and co-culture conditions with NSCG and NFpp10 GBM cells. Phospholipid and lysophospholipid species were quantified by liquid chromatography electrospray ionization tandem mass spectrometry (LC-ESI/MS/MS). Among all lipid species, lysophosphatidylcholines (LPCs) showed marked accumulation in BV2 MG-CM during co-culture of BV2 MG with NSCG (**Figure 4D**) or NFpp10 GBM cells (**Figure 4F**), with the LPC species, LPC 16:0, LPC 18:0, and LPC 18:1 as the most abundant species (**Figures 4E and 4G**). In BV2 MG cells, LPC levels decreased upon co-culture with NSCG cells (**Figure S4B**) and showed no significant change with NFpp10 cells (**Figure S4D**). Notably, CSF1R inhibition (PLX) facilitated increased LPC accumulation in BV2 MG cells in the presence of NSCG (**Figure S4B)** or NFpp10 cells (**Figure S4D**). Like in BV2 MG-CM, LPC 16:0, LPC 18:0, and LPC 18:1 were the predominant LPC species in BV2 cells in both co-culture conditions (**Figures S4C and S4E**).

To validate the effect of PLX on microglial LPC secretion, BV2 MG cells were incubated with a fluorescent LPC derivate (Lyso PC, green) in the presence or absence of PLX. After 24 hours, MG cells had internalized Lyso PC (**Figure 4H**). PLX treatment markedly increased intracellular Lyso PC levels, evident by an increased number of Lyso PC^+^ droplets (**Figure 4H**). Upon subsequent co-culture with NSCG cells for 24 hours, microglia lost most intracellular Lyso PC particles, indicating active lipid release. In contrast, PLX treated BV2 MG cells retained Lyso PC despite the presence of tumor cells (**Figure 4H**). These results demonstrate that CSF1R inhibition promotes intracellular LPC retention in MGs, thereby limiting LPC secretion even under tumor-stimulated conditions (**Figure 4H**).

To establish the *in vivo* relevance of microglia-derived LPCs in GBM invasion, we performed quantitative Mass Spectrometry Imaging (qMSI) on NSCG- and NFpp10-tumor bearing brains (**Figures 4I and 4L, Figures S4F, S4I, S4L and S4O**). K-means lipid clustering revealed distinct spatial lipid distributions across brain and tumor regions of both NSCG- and NFpp10-bearing mice (**Figures 4J and 4M, Figures S4G, S4J, S4M and S4P**). The tumor rim, corresponding to the invasive front where MGs and invading GBM cells interact (NSCG in yellow; NFpp10 in blue/green), displayed a lipid profile distinct from the tumor core (NSCG in green; NFpp10 in orange/red) (**Figures 4J and 4M, Figures S4G, S4J, S4M and S4P**). In NSCG-bearing mice, LPC 16:0, LPC 18:0 and LPC 18:1 were largely absent at the tumor core, whereas LPC 16:0 and 18:0 were enriched at the tumor rim and LPC 18:1 was diffusely distributed throughout the brain (**Figure 4K, Figures S4H and S4K).** In NFpp10-bearing mice, all three LPCs formed clusters at the invasive front, with additional LPC-rich regions detected within the tumor core, and LPC 18:1 also diffusely dispersed (**Figure 4N, Figures S4N and S4Q).** LPC concentrations ranged from 1–4 pmol/mm^2^ in NSCG tumors (**Figure 4N, Figures S4H and S4K**) and 1–6 pmol/mm^2^ in NFpp10 tumors (**Figure 4N, Figures S4N and S4Q**). Laser-capture microdissection of tumor rim and tumor core regions of NSCG and NFpp10 tumors followed by bulk lipidomics, confirmed LPC 16:0, LPC 18:0, and LPC 18:1 as the predominant LPC species in both tumor types (**Figure 4O**). Consistent with qMSI (**Figures 4K and 4N, Figures S4H, S4K, S4N, and S4Q**), LPC levels were higher at the tumor rim than the core in NSCG tumors (**Figure 4O**), while NFpp10 tumors showed no significant regional differences (**Figure 4O**).

These differences likely reflect distinct tumor immune microenvironmental (TIME) landscapes. NSCG tumors are poorly infiltrated by myeloid and immune cells, whereas NFpp10 tumors display dense infiltration by immunosuppressive myeloid cells and dysfunctional T cells^23^. In line with this, consecutive tissue sections to the qMSI samples revealed limited macrophage infiltration in the NSCG tumor core (CD68⁺, in green) (**Figure S4R**), in contrast to the extensive macrophage infiltration observed in NFpp10 tumors (CD68⁺, in red) (**Figure S4S**). This divergence provides a mechanistic explanation for LPC distribution. LPCs are largely confined to the rim of NSCG tumors, consistent with microglia-derived secretion, whereas in NFpp10 tumors, LPCs extend into the core, likely reflecting additional contributions from infiltrating macrophages. Collectively, these findings support MG-secreted lipids as key drivers of glioma invasion, demonstrate that CSF1/CSF1R inhibition suppresses LPC secretion, and identify LPC 16:0, 18:0, and 18:1 as candidate mediators of GBM invasion across GBM types.

### MG-CSF1R and myeloid PI3K inhibition impairs lipid droplet degradation

Next, we sought to gain mechanistic insight into LPC release and retention of GBM-activated MG cells in the absence or presence of PLX. To define the mechanism underlying LPC retention in microglia, we examined how PLX alters lipid handling. Bulk lipidomics of BV2 cells co-cultured with NSCG or NFpp10 tumor cells revealed a substantial accumulation of triglycerides (TGs) upon PLX treatment (**Figure 5A**). TGs, together with cholesteryl esters (CE), form lipid droplets (LDs), organelles that serve as intracellular energy stores of neutral lipids^24^. While CE levels remained unchanged (**Figure S5A**), PLX or IPI treatment induced robust LD accumulation in BV2 MG-cells (**Figure 5B**). This was accompanied by increased expression of *Perilipin-2* (*Plin2*), a key structural protein in the LD membrane, in BV2 cells upon PLX treatment (**Figure S5B**). These effects were recapitulated in freshly isolated murine microglia (**Figure S5C**), and in human HMC3 MG-cells (**Figure S5D**), where LDs similarly accumulated following PLX and/or IPI treatment.

**Figure 5.**
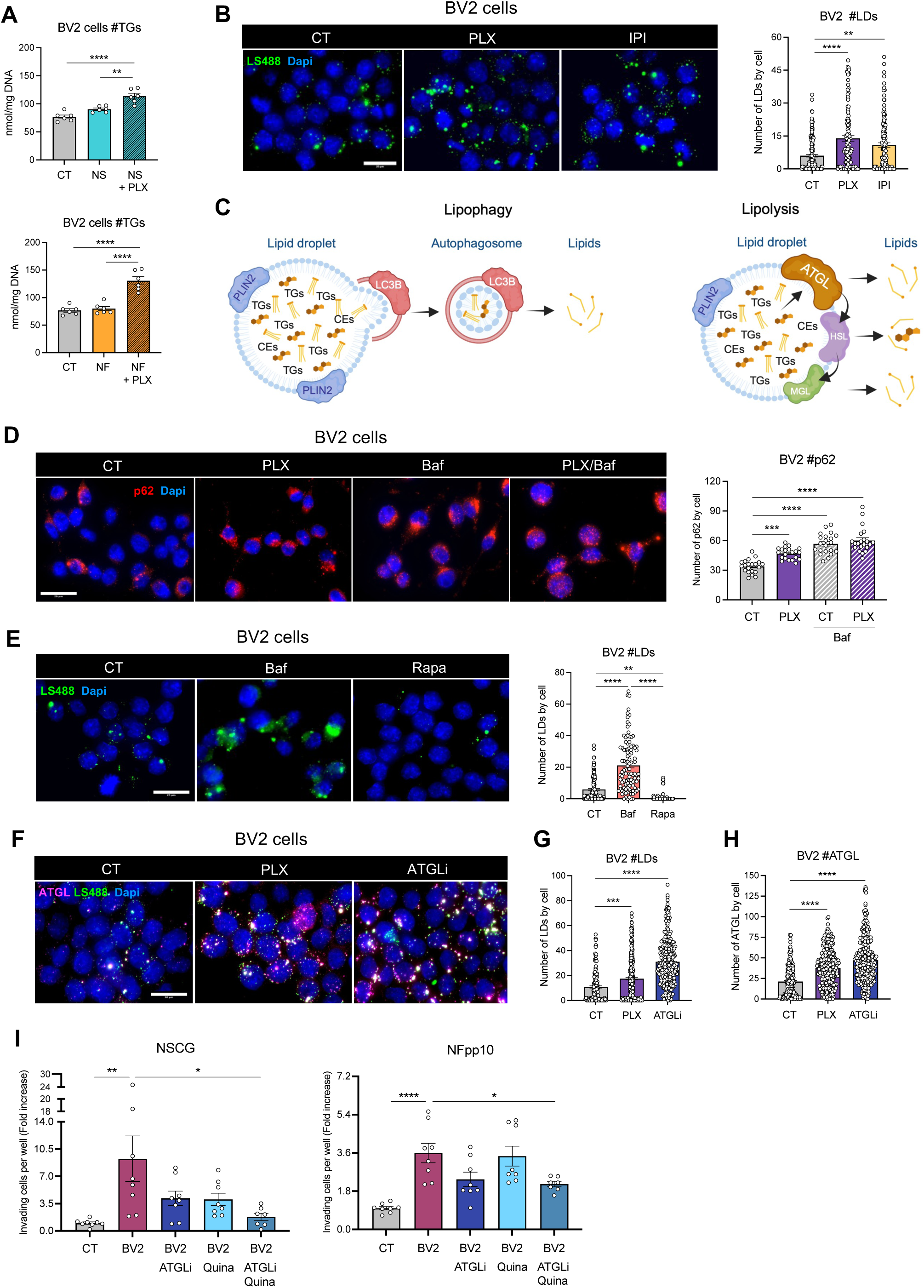
MG-CSF1R and PI3Kγ/δ inhibition blocks lipophagy/lipolysis, leading to lipid droplet accumulation and impaired GBM invasion. (A) Quantification of the sum of TG species in BV2 cells in the absence or presence of NSCG co-culture (upper graph) or NFpp10 co-culture (lower graph) with or without PLX3397. BV2 (NSCG co-culture): n=6, One-way ANOVA, ** p-value=0.0013, **** p-value<0.0001. BV2 (NFpp10 co-culture): n=6, One-way ANOVA, **** p-value<0.0001. (B) Immunofluorescence images and corresponding quantification of BV2 cells (Dapi, blue) and LDs (LS488, green) upon PLX3397 and IPI-145 treatment. Scale bar: 20µm. n=3, One-way ANOVA, ** p-value=0.0010, **** p-value<0.0001. (C) Overview of the two main mechanisms of LD degradation: lipophagy and lipolysis. (D) Immunofluorescence images and corresponding quantification of BV2 cells (Dapi, blue) and p62 (red) upon PLX3397, bafilomycin or both. Scale bar: 20µm. n=3, One-way ANOVA, ** p-value=0.0010, **** p-value<0.0001. (E) Immunofluorescence images and corresponding quantification of BV2 cells (Dapi, blue) and LDs (LS488, green) upon bafilomycin and rapamycin treatment. Scale bar: 20µm. n=3, One-way ANOVA, ** p-value=0.0019, **** p-value<0.0001. (F, G, H) Immunofluorescence images (F) and corresponding quantification of BV2 cells (Dapi, blue), LDs (LS488, green) (G), and ATGL (ATGL, pink) (H) upon PLX3397 and ATGL inhibitor (ATGLi) treatment. Scale bar: 20µm. BV2 – LDs (G): n=7, One-way ANOVA, *** p-value=0.0001,**** p-value<0.0001; BV2 – ATGL (H): n=7, One-way ANOVA, **** p-value<0.0001. (I) Invasion assays of NSCG (left) and NFpp10 (right) cells upon co-culture or not with BV2 untreated or treated with ATGLi, quinacrine (Quina), or both. NSCG: n=4, One-way ANOVA, * p-value=0.0103, ** p-value=0.0026; NFpp10: n=4, One-way ANOVA, * p-value=0.0438, **** p-value<0.0001. All data represent mean ± SEM.

These findings suggest that CSF1R and PI3Kγ/δ inhibition promotes LD accumulation in microglia. Given that LDs are dynamic organelles regulated by degradation pathways, this accumulation suggests impaired turnover. LDs are degraded via two principal mechanisms: lipophagy (autophagic degradation of LDs) and lipolysis (enzymatic breakdown of lipids)^25^ (**Figure 5C**), which we next investigated in the context of PLX treatment.

PLX treatment increased p62 protein levels in both murine BV2 (**Figure 5D**) and human HMC3 microglia (**Figure S5E**) and reduced autophagic flux in HMC3 cells (**Figure S5E**), demonstrating that CSF1R inhibition impairs autophagy. Consistently, pharmacological blockade of autophagy with bafilomycin (**Figure 5E**) or quinacrine (**Figure S5F**), induced similar LD accumulation in both BV2 (**Figure 5E, Figure S5F**) and HMC3 cells (**Figure S5G**). In contrast, autophagy induction with rapamycin, an mTOR inhibitor, reduced LD levels in BV2 MG-cells (**Figure 5E**). These results indicate that CSF1R inhibition suppresses autophagic LD degradation, leading to LD accumulation and limiting the availability of lipids for secretion.

We next assessed the impact of PLX on lipolysis. ATGL (adipose triglyceride lipase), the primary enzyme responsible for mobilizing fatty acids from cellular triglyceride stores, increased in both BV2 and HMC3 MG-cells following PLX treatment, despite persistent LD accumulation (**Figures 5F-H, Figures S5H-J**). A similar phenotype was observed upon pharmacological ATGL inhibition with Atglistatin (ATGLi) (**Figures 5F-H, Figures S5H-J**), which also enhanced LD accumulation. These data suggest that PLX functionally impairs ATGL activity, triggering a compensatory upregulation of ATGL expression without restoring lipolysis to regulate LD accumulation.

We then tested whether inhibiting lipophagy and lipolysis in MG-cells impairs glioma invasion. BV2 MG-cells were pre-treated with atglistatin (ATGLi), quinacrine (Quina), or their combination, followed by *in vitro* invasion assays with NSCG and NFpp10 cells. Inhibition of either pathway alone had no effect; only combined ATGLi and quinacrine treatment reduced glioma invasion (**Figure 5I**), suggestive of the necessity to block both lipophagy and lipolysis to limit tumor invasion. Consistent with this result, combined inhibition increased triglyceride (TG) and cholesteryl ester (CE) levels in BV2 MG-cells (**Figures S5K and S5L**), recapitulating the lipid storage phenotype induced by PLX treatment (**Figure 5A**).

Overall, these findings show that CSF1R and PI3Kγ/δ inhibition drives lipid droplet accumulation in MG cells by suppressing both lipophagy and lipolysis, the two principal pathways of LD degradation.

### Targeting the LPA-LPAR signaling axis impairs GBM invasion

We next tested whether MG-derived LPC 16:0, LPC 18:0, and LPC 18:1 indeed drive migratory and glioma invasion, building on our prior *in vitro* and *in vivo* analyses (**Figures 4D-4G**, **4K, and 4N, Figures S4B-S4E**, **S4H, S4K, S4N, and S4Q**). Among these, only LPC 18:1 increased invasion in NSCG cells (**Figure 6A**), whereas none of the three LPC species affected NFpp10 invasion (**Figure 6B**). Because LPC can be enzymatically converted to lysophosphatidic acid (LPA) by autotaxin (ATX), we next examined *Enpp2* expression (ATX, encoded by *Enpp2*) in NSCG tumor-bearing brains using multiplexed error-robust fluorescence *in situ* hybridization (MERFISH) spatial transcriptomics. *Enpp2* expression was enriched at the NSCG tumor surroundings and overlapped with the expression of MG-specific markers, including *Tmem119* (**Figure 6C**). Transcriptomic data analysis from the TCGA-GBM Pan-Cancer atlas (cBioPortal) of human GBM samples confirmed expression of ENPP2 in microglia (Tmem119^+^, P2RY12^+^) (**Figure 6D**). These findings infer that MG-secreted LPCs can be converted to LPA via ATX at the invasive tumor rim.

**Figure 6.**
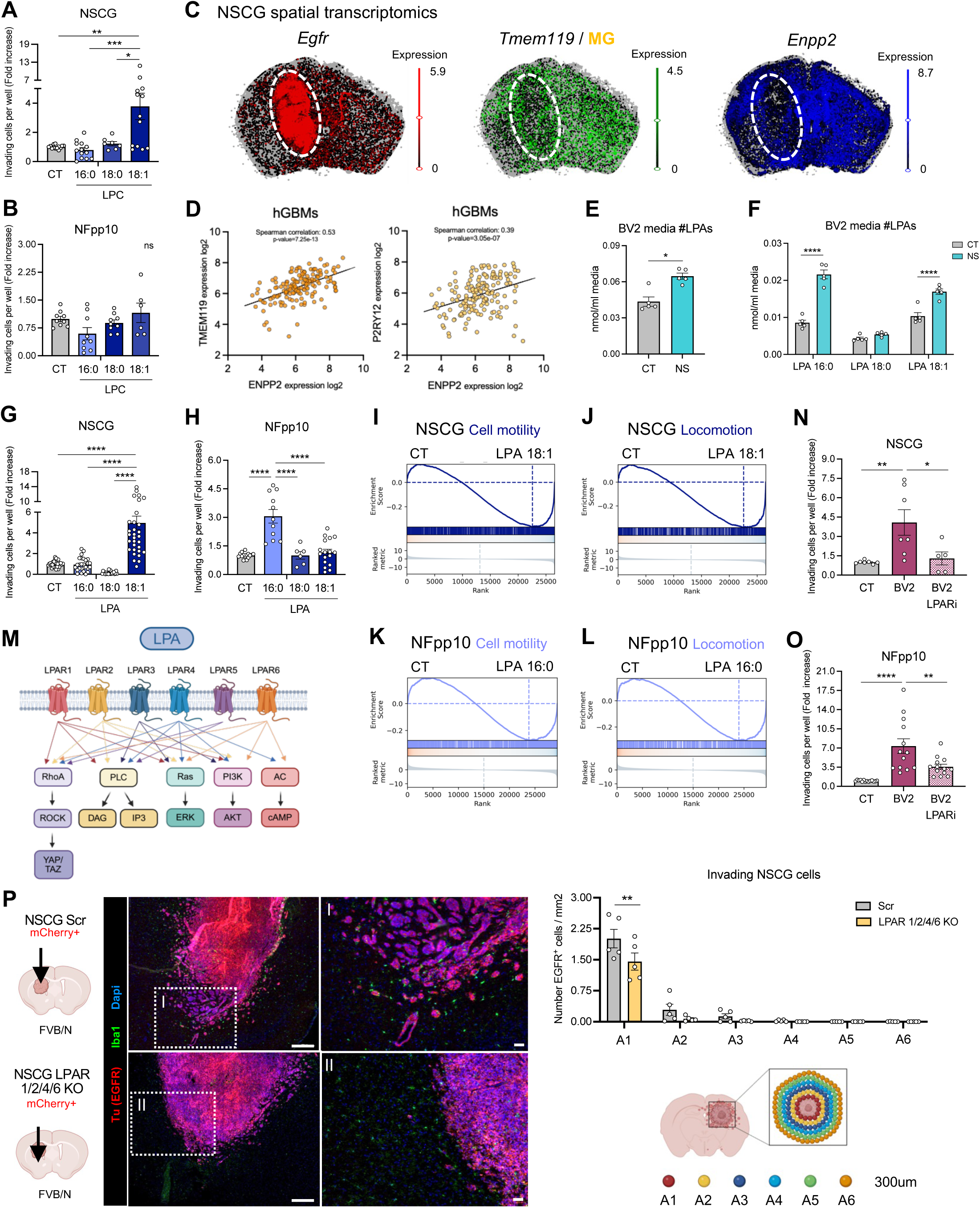
LPA-LPAR signaling promotes GBM invasion. (A) Invasion assays of NSCG cells in the presence or absence of LPC 16:0, LPC 18:0, LPC 18:1. Fold increase. NSCG: n=5, One-way ANOVA, * p-value=0.0190, ** p-value=0.0014, *** p-value=0.0006. (B) Invasion assays of NFpp10 cells in the presence or absence of LPC 16:0, LPC 18:0, LPC 18:1. Fold increase. NFpp10: n=3, One-way ANOVA, non-significant. (C) Spatial transcriptomics analysis of NSCG tumor brains showing *Egfr, Tmem119*, and *Enpp2* expression. (D) Correlation plot of *ENPP2*, *TMEM119*, and *P2RY12* expression in human primary GBM samples. TCGA-Pancancer-cbioportal. Spearman correlation Tmem119/ENPP2: 0.53, p-value=7.25E-13. Spearman correlation P2RY12/ENPP2: 0.39, p-value=3.05E-7. (E) Quantification of the sum of LPA species in BV2 media in the absence or presence of NSCG co-culture. n=5. Mann-Whitney test, * p-value=0.0159. (F) Quantification of the distinct LPA species in BV2 media in the absence or presence of NSCG co-culture. n=5. Two-way ANOVA, ****p-value<0.0001. (G) Invasion assays of NSCG cells in the presence or absence of LPA 16:0, LPA 18:0, LPA 18:1. Fold increase. n=11, One-way ANOVA, **** p-value<0.0001. (H) Invasion assays of NFpp10 cells in the presence or absence of LPA 16:0, LPA 18:0, LPA 18:1. Fold increase. n=6, One-way ANOVA, **** p-value<0.0001. (I) bulkRNAseq leading edge plot of the cell motility GOBP in NSCG cells control and LPA 18:1 treated. p-adj=2.52E-11. NES=1.744043. (J) bulkRNAseq leading edge plot of the locomotion GOBP in NSCG cells control and LPA 18:1 treated. p-adj=1.64E-11. NES=1.759690. (K) bulkRNAseq leading edge plot of the cell motility GOBP in NFpp10 cells control and LPA 16:0 treated. p-adj=0,015153. NES=0,691534. (L) bulkRNAseq leading edge plot of the locomotion GOBP in NFpp10 cells control and LPA 16:0 treated. p-adj= 0.000372. NES=0.941794. (M) General overview of the LPA-LPAR signaling pathways involved in cell migration, differentiation, and proliferation. (N) Invasion assays of NSCG cells untreated or treated with a cocktail of LPAR inhibitors in the presence or absence of BV2 cells. Fold increase. n=3, One-way ANOVA, * p-value=0.0324, ** p-value=0.0098. (O) Invasion assays of NFpp10 cells untreated or treated with a cocktail of LPAR inhibitors in the presence or absence of BV2 cells. Fold increase. n=6, One-way ANOVA, ** p-value=0.0060, **** p-value<0.0001. (P) Representative immunofluorescent images and quantification of invading NSCG control (Scr) and LPAR1/2/4/6 knock-out cells (EGFR^+^, red) and their association with microglial cells (Iba1^+^, green). Scales (upper panels to lower panels): 200µm, 50µm. n=5. One-way ANOVA, ** p-value=0.0014. All data represent mean ± SEM.

Although LPA could not be readily detected by qMSI, dedicated protocol developments in LPA detection^26^ enabled us to quantify LPAs by bulk analysis from BV2 MG-CM under mono- and co-culture conditions with NSCG or NFpp10 cells. Notably, only LPA 16:0, LPA 18:0, and LPA 18:1 were detected and their levels increased in the presence of both NSCG and NFpp10 cells (**Figures 6E and 6F, Figures S6A and S6B**).

We next tested the functional impact of the individual LPA species (LPA 16:0, LPA 18:0, and LPA 18:1) on glioma invasion. Consistent with the LPC results, LPA 18:1 selectively increased invasion in NSCG cells whereas LPA 16:0 or LPA 18:0 had no effect (**Figure 6G**). Interestingly, in NFpp10 cells, LPA 16:0, but not LPA 18:0 or LPA 18:1, induced invasion (**Figure 6H**). Notably, although LPC species did not promote NFpp10 invasion (**Figure 6B**), LPA 16:0 was sufficient to do so (**Figure 6H**).

Finally, bulk RNA-seq of LPA 18:1-treated NSCG cells, and LPA 16:0-treated NFpp10 cells revealed consistent upregulation of cell motility and locomotion pathways in both NSCG and NFpp10 cells, supporting direct and selective LPA signaling in promoting glioma invasiveness (**Figures 6I-6L**).

LPAs are signaling lipids that signal through six G-protein coupled receptors (LPAR1-6) to regulate cell migration, proliferation, survival, and invasion^27^ (**Figure 6M**). To determine whether LPA–LPAR signaling promotes glioma invasion, we pre-treated NSCG and NFpp10 cells with a combination of LPAR inhibitors and assessed invasion in the presence or absence of BV2 MG-cells. LPAR inhibition markedly reduced invasion of both NSCG and NFpp10 cells under co-culture conditions with BV2 MG-cells (**Figures 6N and 6O**), indicating that microglia-driven invasion depends on LPA receptor signaling. To validate the critical role of LPA-LPAR signaling in promoting GBM invasion *in vivo*, we identified Lpar1, Lpar2, Lpar4, and Lpar6 to be preferentially expressed in NSCG cells and generated combined CRISPR/Cas9 knockouts (**Figure S6C**). Consistent with the *in vitro* results, LPAR-deficient NSCG tumors exhibited a significantly reduced invasive capacity compared to controls (**Figure 6P**), establishing a functional requirement for LPA signaling in glioma invasion.

### EGFRvIII^+^ GBM invasion relies on the Yap/Taz signaling pathway

We next sought to define the downstream pathways mediating LPA–LPAR invasion. Single-cell RNAseq analysis revealed enrichment of Yap/Taz signaling in NSCG tumor rim cells compared to tumor core (**Figure 7A**). As LPA-LPAR signaling activates RhoA, a known upstream regulator of Yap/Taz (**Figure 6M**), these data suggest a mechanistic link between microglia-derived LPA and Yap/Taz activation. This enrichment was not observed in NFpp10 tumor rim cells (**Figure 7B**). We validated these findings *in vitro* by assessing canonical Yap target gene expression (*Ankrd*1, *Ctgf*, *Cyr61*) in NSCG and NFpp10 cells co-cultured with BV2 MG-cells. BV2 co-culture increased Yap target gene expression in NSCG cells but not in NFpp10 cells (**Figure 7C**). Consistently, LPA 18:1 selectively induced Yap target gene expression in NSCG cells whereas LPA 16:0 or LPA 18:0 had no effect (**Figure 7D**); none of these LPA species altered Yap target gene expression in NFpp10 cells (**Figure 7E**).

**Figure 7.**
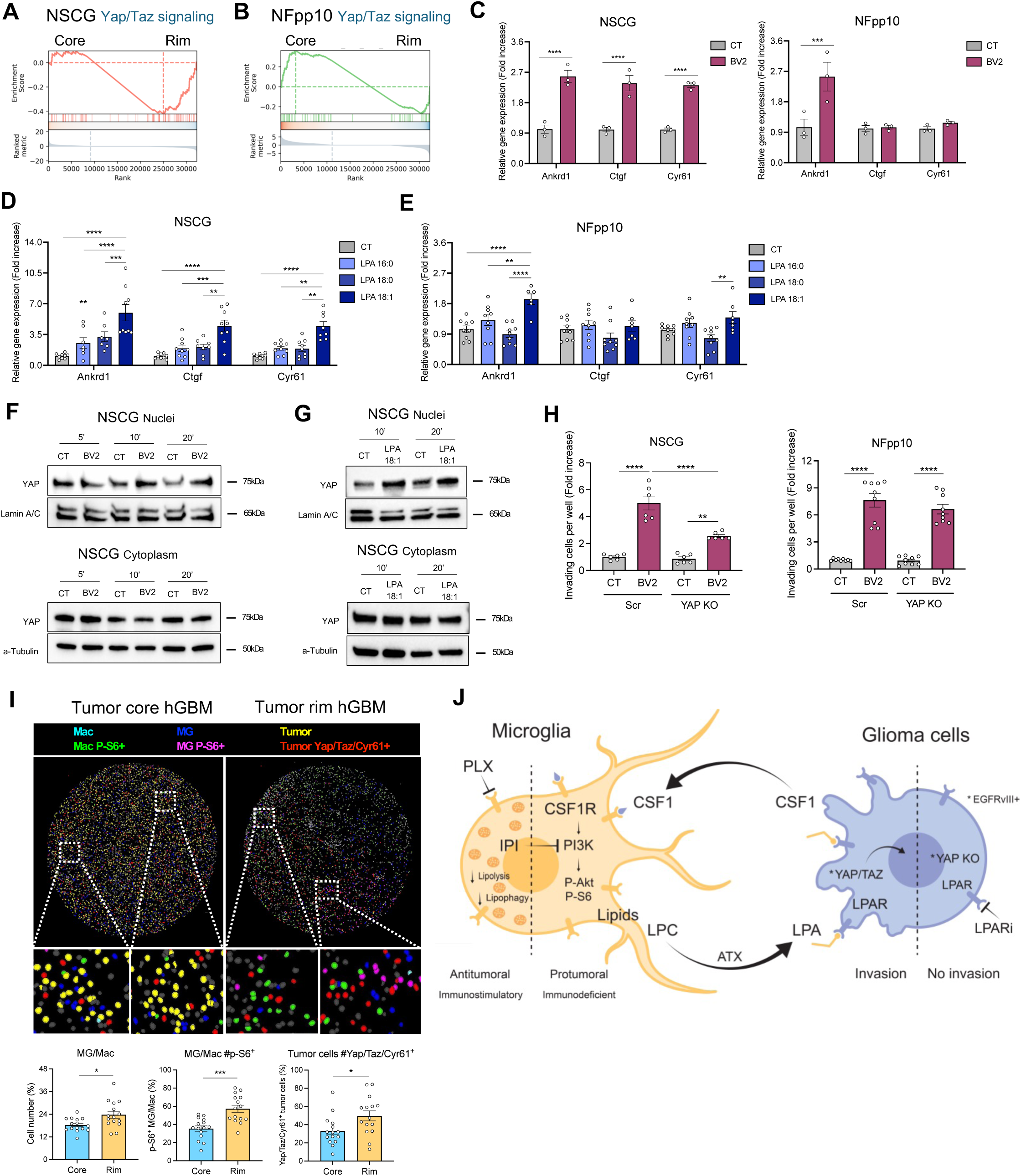
EGFRvIII^+^ GBM invasion relies on the Yap/Taz signaling pathway. (A) scRNA-seq analysis of NSCG tumor rim and core samples focusing on Yap/Taz signaling (Kanai signature). p-adj=4.41E-04. ES= 3.623954. (B) scRNA-seq analysis of NFpp10 tumor rim and core samples focusing on Yap/Taz signaling (Kanai signature). p-adj=6.79E-01. ES= 0.414527. (C) Gene expression of Yap-target genes (*Ankrd1, Ctgf, Cyr61*) in NSCG and NFpp10 cells upon 24 hours of co-culture with BV2 cells. Relative gene expression to S18. Fold increase. NSCG: n=3, 2way ANOVA, **** p-value<0,0001; NFpp10: n=3, 2way ANOVA, Mean -+ SEM, *** p-value=0,0007. (D) Gene expression of Yap-target genes (*Ankrd1, Ctgf, Cyr61*) in NSCG cells upon 24 hours of LPA 16:0, LPA 18:0, and LPA 18:1 treatment. Relative gene expression to S18. Fold increase. n=3. 2way ANOVA, ** p-value=0,0084, *** p-value=0,0007, **** p-value <0,0001. (E) Gene expression of Yap-target genes (*Ankrd1, Ctgf, Cyr61*) in NFpp10 cells upon 24 hours of LPA 16:0, LPA 18:0, and LPA 18:1 treatment. Relative gene expression to S18. Fold increase. n=3. 2way ANOVA, ** p-value=0,0096, *** p-value=0,0007. (F) Representative immunoblots of Yap nuclear and cytoplasmic levels in NSCG cells upon 5 minutes, 10 minutes, 20 minutes of co-culture with BV2 cells. (G) Representative immunoblots of Yap nuclear and cytoplasmic levels in NSCG cells upon 10 minutes and 20 minutes of LPA 18:1 treatment. (H) Invasion assays of NSCG (left) and NFpp10 (right) Scr (control) and YAP knock-out cells in the presence or absence of BV2 cells. Fold increase. NSCG: n=3, One-way ANOVA, ** p-value= 0.0039, **** p-value<0.0001; NFpp10: n=3, One-way ANOVA,**** p-value<0.0001. (I). Multiplex immunohistochemistry and quantification of 18 different GBM patients across 50 TMAs. 15 tumor core TMAs and 15 tumor rim TMAs stratified by tumor cell density. n=30. MG/Mac: Mann-Whitney t test, Mean ± SEM, * p-value=0,0215; MG/Mac P-S6^+^: Mann-Whitney t test, *** p-value=0,0004; Tumor cells Yap/Taz/Cyr61^+^: Mann-Whitney t test, * p-value=0,0164. (J) General overview of the MG-GBM crosstalk. All data represent mean ± SEM.

Similar findings were observed when we treated NSCG cells with BV2-conditioned medium. BV2 MG-CM induced Yap target gene expression selectively in NSCG cells but not in NFpp10 cells (**Figure S7A**). Lipid depletion of BV2 CM abolished this effect, and reconstitution with LPA 18:1, but not LPA 16:0, restored Yap target gene expression in NSCG cells without affecting NFpp10 cells confirming that LPA 18:1 is the active lipid species driving Yap activation in NSCG cells (**Figure S7A**).

Because Yap is transcriptionally active upon nuclear translocation, we quantified nuclear Yap protein levels. BV2 co-culture increased nuclear Yap levels in NSCG cells (**Figure 7F**) as did acute LPA 18:1 stimulation within 10–20 minutes (**Figure 7G**). NFpp10 cells showed no increase in nuclear Yap following BV2 co-culture or LPA 16:0 treatment (**Figure S7B**). Consistent with these findings, Rho–GTP pulldown assays revealed rapid RhoA activation within 5 minutes in NSCG cells after BV2 co-culture or LPA 18:1 treatment (**Figures S7C and S7D**), whereas NFpp10 cells exhibited no detectable Rho activation under identical conditions (**Figures S7C and S7D**).

To validate the functional role of Yap/Taz signaling in glioma invasion, particularly in EGFRvIII⁺ GBMs, we generated CRISPR/Cas9 knockouts of *Yap* and its partner *Taz* in NSCG and NFpp10 cells. Loss of Yap/Taz significantly reduced invasion of NSCG cells, both at baseline and during co-culture with BV2 microglia (**Figure 7D, Figure S7E**) and diminished the pro-invasive effect of LPA 18:1 (**Figure S7F**). In contrast, Yap/Taz deletion had no effect on NFpp10 invasion (**Figure 7H, Figure S7E**).

Collectively, these findings demonstrate that microglia-derived LPA 18:1 drives NSCG invasion through a RhoA-Yap/Taz signaling axis. NFpp10 cells, in contrast, invade through a Yap/Taz independent mechanism, underscoring subtype-specific wiring of LPA signaling.

To validate these findings in human GBM, we performed multiplex immunohistochemistry (MILAN) on tissue microarrays (TMA) encompassing 50 GBM regions from 18 patients. TMAs were stratified by tumor cell density, with the 15 highest density regions defined as core, and the 15 lowest as rim (**Figure S7G**). Tumor rim regions exhibited increased microglia/macrophage infiltration compared to tumor core (**Figure 7I**), resulting in a reduced tumor cell-to-microglia/macrophage ratio (**Figure S7H**). Consistently, tumor cell density inversely correlated with microglia/macrophage abundance (**Figure S7I**). GBM-associated microglia/macrophages showed elevated PI3K activation (via P-S6 levels) at the tumor rim (**Figure 7I**), recapitulating our observations in murine GBM (**Figures 1A–1D**). Concurrently, tumor cells at the rim displayed increased expression of YAP, TAZ and CYR61, supporting their activation of a pro-invasive transcriptional program (**Figure 7I**). Together, these data indicate that microglia–tumor cell interactions at the invasive front are conserved in human GBM and are associated with activation of both microglial signaling and tumor cell YAP/TAZ activity.

## Discussion

The infiltrative growth of GBM constrains surgical resection, the strongest determinant of patient survival, underscoring the need to define and target mechanisms that drive tumor invasion. Here, we delineate a paracrine signaling axis in which rim-localized GBM cells activate CSF1R–PI3K signaling in microglia, triggering the release of bioactive lipids, including LPCs. These LPCs are further converted by ATX into LPA species that engage LPA receptors on GBM cells to promote migration. Notably, although microglia secrete a diverse repertoire of LPC species, the migratory response of GBM cells is genotype dependent. In EGFRvIII⁺ tumors, lysophosphatidic acid (LPA 18:1)–LPAR signaling activates a RhoA–Yap/Taz cascade, driving nuclear translocation of Yap/Taz and the induction of pro-invasive transcriptional programs. By contrast, LPA 16:0, but not LPA 18:1, promotes invasion in NF1/p53/PTEN-deficient GBM, and is Yap/Taz-independent. Together, these findings establish a context-dependent lipid signaling axis that couples microglial activation to glioma invasion *in vitro* and *in vivo* (**Figure 7J**).

Beyond invasion, YAP/TAZ activation has been implicated in GBM proliferation, survival, stemness and therapeutic resistance^28–31^, and more broadly in tumor progression, survival, epithelial-mesenchymal transition (EMT), and recurrence across multiple cancer types including non-small cell lung cancer (NSCLC)^32^, hepatocellular carcinoma^33,34^, breast^35^, colorectal^36^, and ovarian cancers^37^. So far, paracrine interactions between GBM and CSF1R-activated microglia have predominantly been attributed to growth factors and chemokines such as EGF and TGFβ, Amphiregulin (AREG) and plexin-B2-mediated signaling^38–42^.

By contrast, the contribution of lipid-mediated signaling to GBM invasion remains poorly defined. Our results position lipid signaling as a central component/node of the GBM-MG crosstalk *in vitro* and *in vivo*, with microglia serving as a major source of LPC and LPA within the central nervous system (CNS)^43–45^. Our findings also align with recent studies were elevated LPA levels, increased LPA receptor (LPAR), and autotaxin (ATX) expression have been associated with tumor progression and metastasis across multiple cancer types, including GBM^44–47^, colon cancer^48,49^, breast cancer^50–52^, ovarian cancer^53,54^, pancreatic cancer^55^, and fibrosarcoma^56^. Notably, MG-secreted LPAs have also been linked to various inflammatory conditions in the brain, leading to demyelination, synaptic dysfunction, neurotoxicity, blood-brain barrier disruption, and contributing to secondary injuries in several neuroinflammatory and neurodegenerative diseases, including multiple sclerosis, Alzheimer’s, ischemic stroke, and traumatic brain injuries^43,57–60^.

Finally, our data suggest that therapeutic disruption of the pro-invasive GBM–microglia circuit can restrain glioma invasion. We identify multiple intervention points along this axis that block glioma invasion. Disruption of CSF1R–PI3K signaling in microglia, inhibition of lipid mobilization through suppression of lipophagy and lipolysis, blockade of LPA–LPAR signaling in glioma cells, or inhibition of YAP/TAZ activity in EGFRvIII⁺ GBM each attenuates glioma cell migration, indicating that this paracrine circuit contributes functionally to tumor invasion and is amenable to therapeutic targeting *in vivo*. We found that pharmacological inhibition of CSF1R reprograms microglia toward a less tumor-permissive state, concomitant with reduced LPC secretion and diminished support of tumor invasion, and it is accompanied by intracellular lipid accumulation consistent with reduced lipophagy and lipolysis. Previous studies targeting CSF1R have reported reduced tumor growth, enhanced survival, and improved response to temozolomide in proneural GBM mouse models^61–65^. However, these effects are context-dependent, as CSF1R inhibition shows limited survival benefits across other GBM subtypes^62,66^, including our models, and the CSF1Ri PLX3397 failed to demonstrate clinical benefit in early-phase trials^67^. These observations suggest that targeting the microglial compartment alone is likely insufficient to improve survival of patients with established brain tumors. Instead, our findings raise the possibility that modulating microglial cells before surgical intervention could constrain tumor infiltration and promote a more circumscribed growth pattern, thereby improving resectability of GBMs.

Downstream of microglial lipid release, the ATX–LPA–LPAR signaling axis represents a complementary therapeutic entry point. Although inhibitors targeting ATX or individual LPA receptors have advanced into clinical trials, primarily in fibrotic diseases, their efficacy in cancer remains to be established. For example, LPAR1 antagonists such as BMS-986278 and HZN-825 have progressed to late-stage clinical evaluation in fibrotic disorders but have provided limited or no efficacy in some indications, underscoring the challenges of targeting this pathway through single-receptor inhibition. The redundancy of LPA receptor subtypes and the complexity of LPA signaling likely constrain the effectiveness of such approaches, suggesting that broader pathway inhibition may be required. At the tumor cell level, YAP/TAZ emerges as a key downstream effector of lipid-driven invasion, particularly in EGFRvIII⁺ GBMs. Pharmacological inhibitors that disrupt YAP/TAZ–TEAD interactions are currently being evaluated in early-phase clinical trials for solid tumors, where they have shown preliminary signs of disease stabilization and partial responses in selected patients^68^. These agents provide a potential strategy to interfere with invasion downstream of microenvironmental cues.

Collectively, our findings define a previously unrecognized lipid-mediated crosstalk that links microglial activation to glioma invasion and highlight multiple points of therapeutic vulnerability. Targeting this axis offers a strategy to constrain glioma invasion, enhance surgical resection, and improve clinical outcome.

## Methods

### Murine tumor models

NSCG-mCherry^+^ GBM cells originated from FVB/N CDKN2A-deficient murine neural stem cells (NSCs) with ectopic EGFRvIII expression. NFpp10-GFP^+^ GBM cells were derived from C57BL/6 NSCs with Trp53, Nf1, and Pten-knockdown.

For *in vitro* assays, NSCG cells were cultured in MEM-alpha media (Gibco, 22561-021) supplemented with 5% FBS, 1.2% Glucose 45% (Sigma, G8769), 1% Hepes buffer (Gibco, 15630080), and 1% penicillin-streptomycin (Thermo Fisher, 15140122). NFpp10 cells were cultured in Advanced DMEM/F12 media (Gibco, 12634-010) supplemented with 5% FBS, 1% N2-supplement (Gibco, 17502-048), 1% L-glutamine (Thermo Fisher, 25030024), 20ng/ml FGF-2 (Preprotech, 100-18b), 20 ng/ml EGF (Proprotech, AF-100-15), 50 ug/ml Heparin (Sigma, H3393), and 1% penicillin-streptomycin (Thermo Fisher, 15140122).

For *in vivo* assays, NSCG and NFpp10 cells were cultured as spheres in DMEM/F12 1:1 media (Gibco, 11320-033) supplemented with, 0,5% Hepes (Gibco, 1563-056), 1% B27 supplement without vitamin A (Gibco, 12587-010), 10ng/ml FGF-2 (Preprotech, 100-18b), 10ng/ml EGF (Proprotech, AF-100-15), and 1% penicillin-streptomycin (Thermo Fisher, 15140122). Cell lines were passaged following dissociation using TrypLE (Thermofisher, 12605010) upon reaching 70% confluency.

### Patient-derived cell lines (PDCLs)

EGFR amplified/mutated^+^ cells: LBT012 LamB-GFP^+^ GBM cells were isolated from the tumor core of a GBM patient, and LBT090 Lami-GFP^+^ GBM cells were isolated from the tumor rim of a GBM patient.

EGFR wildtype cells: LBT065 LamB-GFP^+^ and LBT123 LamB-GFP^+^ GBM cells originated from the tumor core of GBM patients, and LBT007 Lami-GFP^+^ and LBT059 Lami GBM cells were isolated from the tumor rim of GBM patients.

For *in vitro* and *in vivo* assays, human GBM cell lines were cultured in DMEM F-12 glutamine-containing media (Life technologies BIPP, 11320074) supplemented with 50ug/ml heparin (Sigma, H3393), 20ng/ml FGF-2 (Stem Cell technologies, 78003.2), 20ng/ml EGF (Stem Cell technologies # 78006.2), 1% antibiotic antimycotic (Life technologies, 15240-062), and a cocktail of additives formed by 30% glucose (Sigma, G7021), Hepes (Gibco, 1563-056), L-glutamine (Thermo Fisher, 25030024), bovine transferrin (Life technologies, 11108016), insulin (Sigma, I2643), putrescine (Sigma, P7505), sodium selenite (Sigma, S9133) and progesterone (Sigma, P6149).

Prior seeding, flasks were coated with 0,5% laminin (Sigma, L2020) in D-PBS (Thermo Fisher, 14190250) during an hour at 37 C.

Cell lines were passaged following dissociation using Accutase (Gibco, A1110501) upon reaching 80% confluency.

### Microglia models

BV2 murine microglial cells were cultured in DMEM high glucose (Gibco, 41965062) supplemented with 10% FBS, 1% L-glutamine (Thermo Fisher, 25030024), and 1% penicillin-streptomycin (Thermo Fisher, 15140122). HMC3 human microglial cells were cultured in DMEM/F12 (1:1) (Gibco, 11320033) supplemented with 10% FBS, 1% L-glutamine (Thermo Fisher, 25030024), and 1% penicillin-streptomycin (Thermo Fisher, 15140122).

BV2 and HMC3 cell lines were passaged following dissociation using TrypLE (Thermofisher, 12605010) upon reaching 70% confluency.

Fresh-isolated murine microglial cells were obtained from FACS-sorted mouse brains. Briefly, mouse brains were mechanically digested in HBSS media (Gibco, 24020091) supplemented with 15mM Hepes (Gibco, 1563-056), and 0,5% Glucose 45% (Sigma, G8769), using the gentleMACS Octo Dissociator (Miltenyi, 2201612). Samples were filtered, washed with 2,5% FBS and 2.5mM EDTA (VWR, E177) in D-PBS (Thermo Fisher, 14190250), and centrifuged 5 min at 1500 rpm. Cell pellet was resuspended in Percoll 38% (Sigma, P1644). Sample was centrifuged at 2000 rpm during 20 min at 20°C. Cell pellet was washed with 5% FBS in D-PBS (Thermo Fisher, 14190250) and centrifuged 5 min at 1500 rpm. Cell pellet was resuspended in ACK Lysis Buffer (15 mM NH4Cl, 10 mM KHCO3, 0.1 mM Na2EDTA) and incubated for 5 min at room temperature. Sample was washed with 5% FBS in D-PBS (Thermo Fisher, 14190250) and centrifuged 5 min at 1500 rpm. Remaining cell pellet was stained for CD45 BV605 (Biolegend, 103139), CD11b APC (Thermo Scientific,17-01112-83), and CX3CR1 FITC (Biolegend, 149019), and microglial cells were FACS-sorted for CD45 intermediate/CD11b^+^/CX3CR1^+^ cell status.

Isolated-microglial cells were directly sorted in 100% FBS and plated in DMEM/F12 (1:1) media (Gibco, 11320033) supplemented with 20% FBS, 1% N2-supplement (Gibco, 17502-048), and 1% penicillin-streptomycin (Thermo Fisher, 15140122). They were maintained for maximum 7 days *in vitro*.

### Ethics statement

All experiments were performed with ethical approval from the Institutional Animal Care and Research Advisory Committee of KU Leuven (ECD 190/2021) and were performed following the institutional and national guidelines and regulations. Human studies were performed under protocols S-68937 and S-62248.

### Animal studies

6-8 weeks old FVB/N, C57Bl/6, and Rag2 female and male mice were anesthetized using isofluorane and placed on a stereotaxic machine. Tumor cells (5×10^3^ NSCG-mCherry^+^, 2×10^4^ NFpp10-GFP^+^, 5×10^4^ NSCG-mCherry^+^ Scr (control), 5×10^4^ NSCG-mCherry^+^ LPAR1/2/4/6 knock-out, 1×10^6^ LBT012 LamB-GFP^+^, 1×10^6^ LBT090 Lami-GFP^+^, 1×10^6^ LBT065 LamB-GFP^+^, or 1×10^6^ LBT007 Lami-GFP^+^) were injected into the right striatum and grown for 14 days (NSCG, NFpp10 tumors) or 4-5 months (LBT012 LamB, LBT090 Lami, LBT065 LamB, LBT007 Lami tumors). PLX treatment started 1 day after tumor cell injection and consisted of PLX5622 containing food pellets versus control food pellets. Upon sacrifice, brains were collected for analysis.

### Immunofluorescence: tissue and cell stainings

<u>-Tissue staining:</u> mouse tumors were fixed in 2% PFA overnight at 4°C, infiltrated with 30% sucrose, and embedded and frozen in OCT compound. Ten μm thick frozen sections were cut using a Cryostar NX70 cryostat (Thermofisher) and collected on Superfrost Plus slides (Fisher Scientific, 22-037-246). Tissue sections were permeabilized and blocked for 1 hour at room temperature with 5% donkey serum (Jackson ImmunoResearch, 017-000-584 121), 0.3% Triton X-100 (Sigma, T8787), and 2% BSA (Sigma, 10735094001). Primary antibodies listed in Table 1 were diluted in respective blocking buffers and incubated overnight at 4°C. Alexa Fluor-conjugated secondary antibodies listed in Table 2 were diluted 1:800 in respective blocking buffers and incubated 1 hour at room temperature. DAPI (4’,6-diamidino-2-phenylindole) (Sigma, D9564) was used for the nuclear counterstain and DAKO fluorescence mounting medium (Agilent, S302380-2) was used for mounting of samples. Images were captured using an AxioCam MRM camera (Zeiss) with objectives of 10X, 20X, 40X magnifications and Axioscan Z1 (Zeiss) at 20X magnification, and analyzed using both Axiovision software (Zeiss) and FIJI software.

**Table 1:**
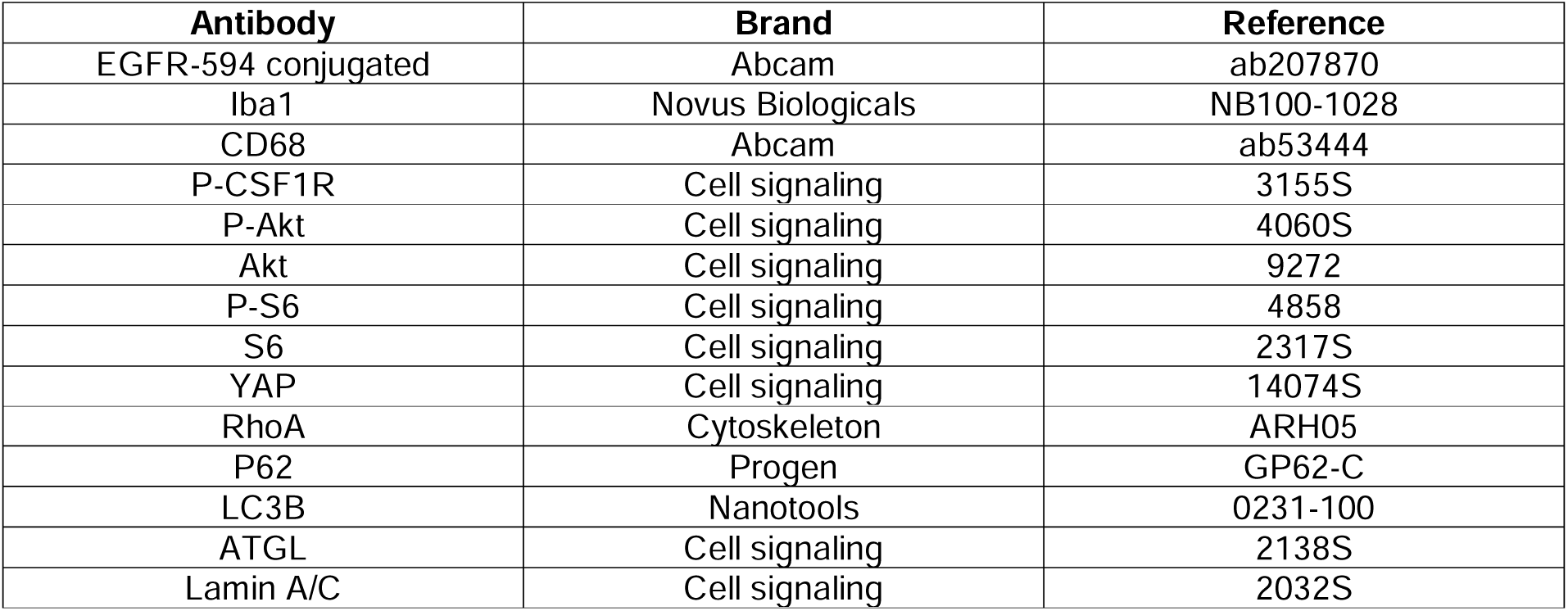
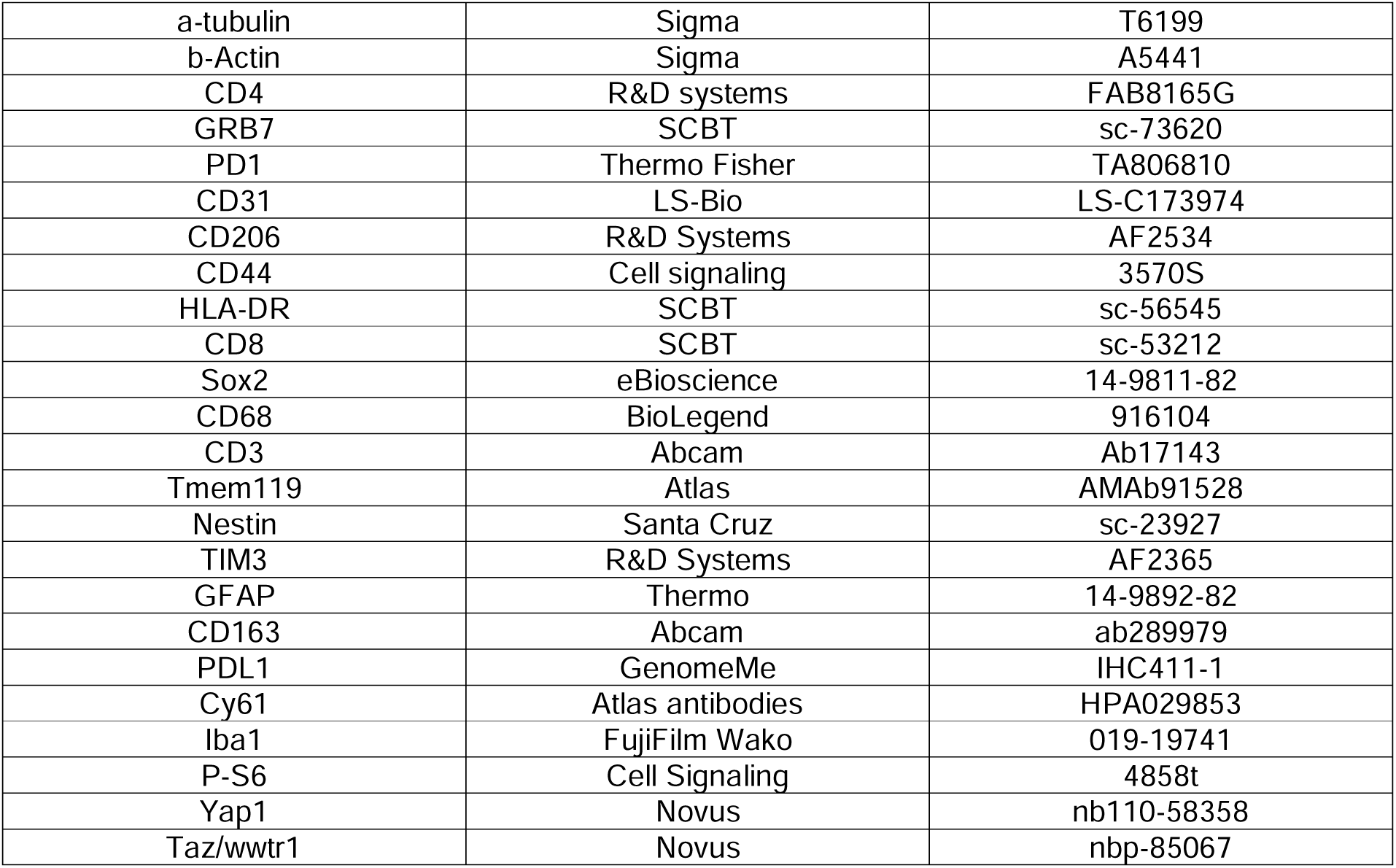
Primary antibodies.

**Table 2:** Secondary antibodies.

| Antibody | Brand | Reference |
| --- | --- | --- |
| Donkey a-rat AF594 | Invitrogen | A21209 |
| Donkey a-goat AF647 | Invitrogen | A21447 |
| Donkey a-mouse AF488 | Invitrogen | A32766 |
| Donkey a-rat AF488 | Life Technologies | A21208 |
| Donkey a-rabbit AF568 | Invitrogen | A10042 |
| Donkey a-rabbit AF647 | Invitrogen | A32795 |
| Goat anti-mouse HRP | SouthernBiotech | 1010-05 |
| Goat anti-rabbit HRP | SouthernBiotech | 4050-05 |
| Goat a-mouse AF488 | Jackson Immuno Research Laboratories | 115-545-206 |
| Goat a-Mouse AF488 | Jackson Immuno Research Laboratories | 115-545-207 |
| Donkey a-rabbit AF647 | Jackson Immuno Research Laboratories | 711-605-152 |
| Goat a-mouse AF555 | Invitrogen | A21127 |
| Goat a-rat AF555 | Invitrogen | 21434 |

<u>-Cell staining:</u> cells were cultured on glass coverslips and fixed using 4% PFA for 30 minutes at 37°C. Coverslips were first permeabilized and blocked with 0,1% Triton X-100 (Sigma, T8787) containing 5% donkey serum (Jackson ImmunoResearch, 017-000-584 121). Primary antibodies listed in Table 1 were diluted in respective blocking buffers and incubated overnight at 4°C. Alexa Fluor-conjugated secondary antibodies listed in Table 2 were diluted 1:800 in respective blocking buffers and incubated 1 hour at room temperature. For lipid spot 488 (dye for neutral lipids), cells were directly incubated 45 minutes at room temperature at 1:1000 dilution in PBS. DAPI (Sigma, D9564) was used for the nuclear counterstain and DAKO fluorescence mounting medium (Agilent, S302380-2) was used for mounting of coverslips. Images were captured using an AxioCam MRM camera (Zeiss) with objectives of 40X, 63X magnifications and analyzed using both Axiovision software (Zeiss) and FIJI software.

### Invasion assays

1,25×10^5^ BV2 cells or 1×10^5^ HMC3 cells were plated on 24-well plates (VWR, 734-1605) in DMEM 10% FBS. When necessary BV2 and HMC3 cells were treated with 2,5µM PLX3397 (MedChemExpress, HY-16749) or 2,5µM PLX5622 (MedChemExpress, HY-114153), and 100nM IPI-145 (LC Laboratories, D-8147).

<u>-For conditioned-media (CM) invasion assays:</u> BV2 and tumor cells were co-cultured during 72 hours in DMEM 10% FBS. After 72 hours, CM was collected, centrifuged, and filtered to remove any possible cell debris and used directly in invasion assays. When necessary, Lipid Removal Agent (LRA) (Sigma, 13358-U) was used to remove the lipids in the CM as indicated in the product protocol prior usage in invasion assays.

<u>-For LPC/LPA treatments:</u> 50µM of LPC 16:0 (Avanti Polar Lipids, 855675C), 50µM LPC 18:0 (Avanti Polar Lipids, 855775C), 50µM LPC 18:1 (Avanti Polar Lipids, 845875C), 50µM LPA 16:0 (Sanbio, 10010093-5), 50µM LPA 18:0 (Sanbio, 10010164-5) and/or 50µM LPA18:1 (Sanbio, 10010093-5) were incubated overnight in 10% FBS DMEM at 4°C in a steering wheel. After overnight incubation, LPC/LPA-enriched media was placed in the invasion assay.

<u>-For quinacrine and ATGL inhibitor (Atglistatin) invasion assays:</u> microglial cells were pre-treated for 48 hours with either 0.5µM quinacrine (Sigma, Q3251), 20µM atglistatin (Med Chem, HY-15859) or both in combination before invasion assay. In addition, quinacrine and atglistatin were also added with the microglial cells during the invasion assay.

Afterward, 0.2mg/ml Matrigel (Corning, 734-1100) was placed in 24-well transwells (Sarstedt, 833932800) and incubated for 30 minutes at 37°C. Matrigel was then removed and 5×10^4^ tumor cells (NSCG wild-type, NSCG CRISPR/Cas9 YAP knock-out, NSCG CRISPR/Cas9 YAP/TAZ knock-out, NFpp10 wild-type, NFpp10 CRISPR/Cas9 YAP knock-out, NFpp10 CRISPR/Cas9 YAP/TAZ knock-out, or PDCLs) in 0% FBS DMEM were plated on the transwells.

For LPAR inhibitor invasion assays: tumor cells were pre-treated for 48 hours with an LPAR inhibitor cocktail constituted by 50µM LPAR1/3/5 inhibitor (Bio-Techne,H2L5765834) and 50µM LPAR2 inhibitor (Sanbio, H2L5186303) before the invasion assay. In addition, the LPAR inhibitor cocktail was also added together with the tumor cells during the invasion assay.

After overnight incubation at 37°C, tumor cells were removed, and transwells were fixed with 100% methanol (VWR, 20847.32) and stained using 0.5% Crystal violet (Sigma, V5265) in 25% methanol. Images were taken using a bright field Axiovert S100 microscope with objective of 10X magnification and analyzed using FIJI software.

### Cell assays

<u>-For Lyso PC experiments:</u> 2,5×10^4^ BV2 cells were plated in coverslips in their own media in 6-well plates (VWR, 734-1599) and 2×10^5^ NSCG cells were plated in their own media in 6-well transwells (VWR, 734-2720). After 6 hours, 2.5µM PLX3397 was added on BV2 cells when needed. Next day, BV2 cells were loaded either with 10µM of TopFluor Lyso PC <u>(Avanti Polar Lipids, 810284P)</u> in combination or not with 2.5µM PLX3397. 24 hours later, BV2 coverslips were washed 3x with D-PBS (Thermo Fisher, 14190250) and some coverslips were fixed and stained with DAPI as indicated in the “**Immunofluorescence: tissue and cell stainings**” method. In addition, the transwells containing NSCG cells were added as co-culture with the remaining BV2 coverslips. 24 hours after co-culture, BV2 coverslips were washed, fixed, and stained with DAPI as indicated before.

<u>-For lipid droplet accumulation experiments:</u> 1×10^4^ BV2 cells, 2×10^4^ HMC3 or 8×10^4^ fresh-isolated microglial cells were plated in coverslips in their own media in 12-well plates (VWR, 734-1598). Next day, BV2, HMC3 or freshly isolated microglial cells were treated for 48 hours either with 2.5 µM PLX3397 (MedChemExpress, HY-16749), 2.5 µM PLX5622 (MedChemExpress, HY-114153), 100nM IPI-145 (LC Laboratories, D-8147), 0.5µM quinacrine (Sigma, Q3251), 2.5µM rapamycin (Selleck chemicals, S1039), or 20µM atglistatin (Med Chem, HY-15859). Exceptionally, BV2 and HMC3 cells were treated only for 6 hours with 50nM bafilomycin (Cayman Chemical, 11038). Coverslips were fixed and stained as indicated in the “<u>Immunofluorescence: tissue and cell</u> <u>stainings</u>” method.

<u>-For Yap-target genes expression experiments:</u> 1.5×10^5^ NSCG, and 1×10^5^ NFpp10 cells were plated in 6-well plates (VWR, 734-1599) in their own media.

For BV2 co-culture experiments, 2.5×10^4^ BV2 cells were plated in 6-well transwells (VWR, 734-2720). After 48 hours, tumor cells and BV2 cells were co-cultured together in 10% FBS DMEM for 24 hours. Cell samples were recollected afterwards for RNA extraction.

For LPA-treatment experiments, 50µM LPA 16:0 (Sanbio, 10010093-5), 50µM LPA 18:0 (Sanbio, 10010164-5) and/or 50µM LPA18:1 (Sanbio, 10010093-5) were incubated overnight in 10% FBS DMEM at 4°C in a steering wheel. After overnight incubation, NSCG and NFpp10 cells were treated with LPA-enriched media for 24 hours.

For conditioned-media (CM) experiments with or without lipids: BV2 and tumor cells were co-cultured during 72 hours in DMEM 10% FBS. After 72 hours, CM was recollected, centrifuged, and filtered to remove any possible cell debris and used directly on the NSCG and NFpp10 cells during 24 hours treatment. When necessary, Lipid Removal Agent (LRA) (Sigma, 13358-U) was used to remove the lipids in the CM as indicated in the product protocol prior usage on the tumor cells. In addition, to rescue the effect of the lipid removal agent, 50µM LPA 18:1 (Sanbio, 10010093-5) and/or 50µM LPA 16:0 (Sanbio, 10010093-5) were incubated overnight in CM + LRA at 4°C in a steering wheel prior treatment to the NSCG and NFpp10 cells for 24 hours.

Cell samples were recollected afterwards for RNA extraction.

### RNA Extraction and Quantitative Reverse-Transcription PCR

Total RNA was extracted from cells using the Pure Link RNA mini kit (Thermo Fisher Scientific, 12183018A) following manufacturer instructions and quantified by Nanodrop. cDNA was reversely transcribed from 1μg of RNA using the iScript Reverse transcription kit (Thermo, 18080-051) according to the manufacturer’s recommendations. Real-time quantitative PCR (RT-qPCR) was performed using the Power Up SyberGreen Master Kit (Thermo Fisher Scientific, A25742). The master mix was prepared according to the manufacturer’s instructions. qPCR was performed on a Biorad CFX96 Real-time system (C1000 Touch thermal cycler) utilizing specific primers at a concentration of 10x. Primers used were obtained from Primer Bank (Harvard University). Primer sequences used in this study were specified in Table 3. Data analysis was performed using the Biorad CFX manager software. CT values were detected for each sample and normalized to the house-keeping gene ribosomal protein S18.

**Table 3:**
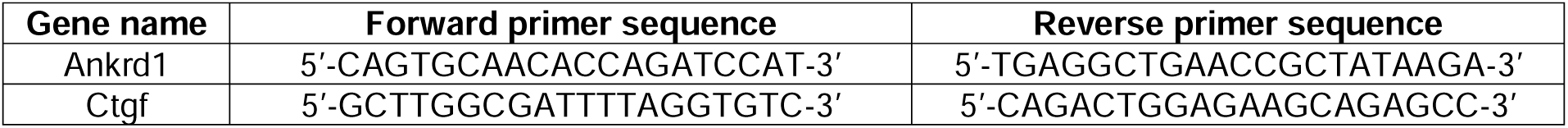
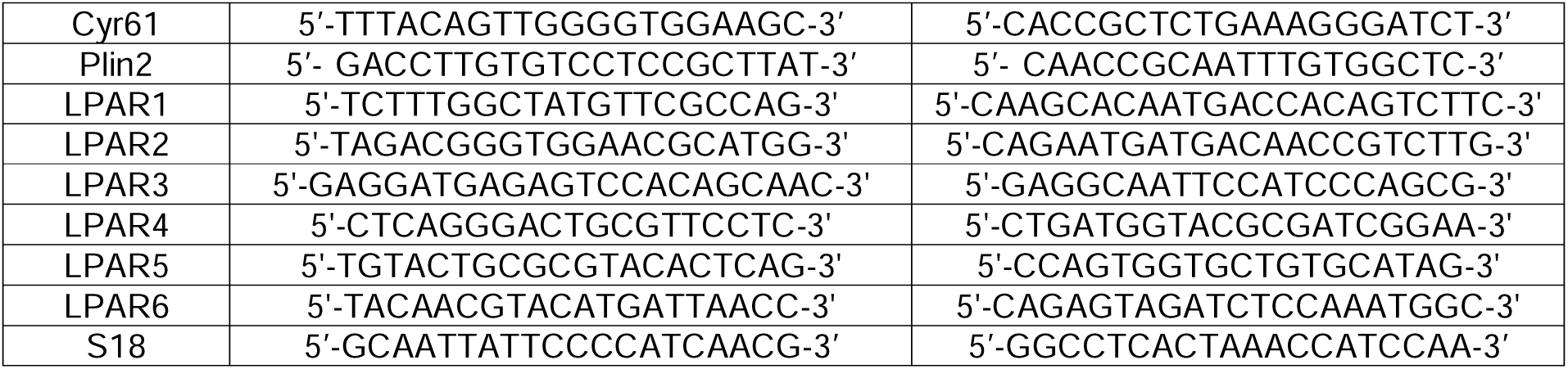
Primer pairs.

**Table 4:**
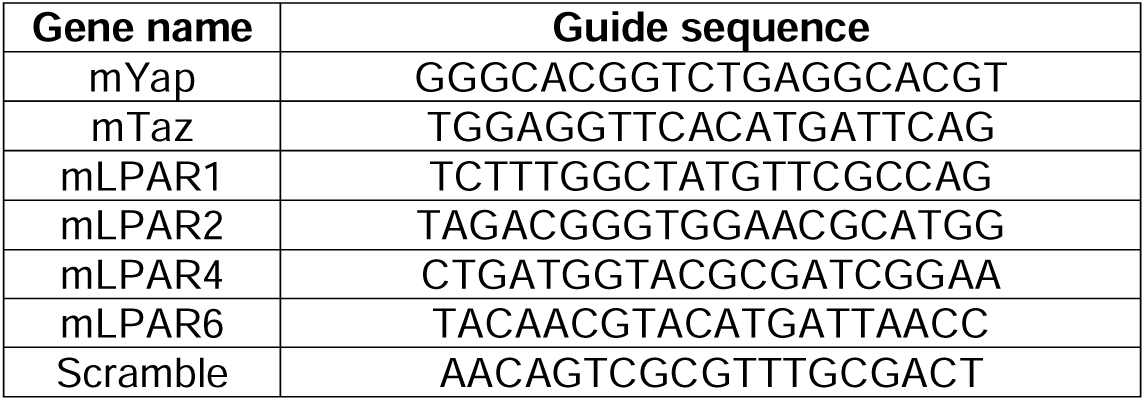
Guide sequences.

### Western blotting

Cells were washed in ice-cold D-PBS (Thermo Fisher, 14190250) and lysed using RIPA buffer (Thermo Scientific, 89901) containing 1:100 phosphatase/protease inhibitor cocktail (Thermo Scientific, 78441). Samples were rotated overnight at 4°C and spin at maximum speed for 15 minutes at 4°C. The supernatants of the samples were collected, and protein concentration was determined by the BCA protein assay kit (Thermo Fisher, 23225). Equal amounts of sample were combined with Laemmli buffer (Biorad, 1610747), denatured for 5 minutes at 95°C, and subjected to SDS/PAGE separation using 4%–20% TGX Precast Gels (Biorad, 4561094), and transferred to PVDF membranes (Biorad, 1704156). Membranes were blocked for 1 hour at room temperature in 5% BSA (Sigma, 10735094001), 0,1% Tween-20 (Sigma, P9416) in D-PBS (Thermo Fisher, 14190250). Incubation with first antibodies (Table 1) was done overnight at 4°C followed by 1 hour incubation at room temperature with secondary antibodies (Table 2). Proteins were visualized using ECL western blot reagents (Biorad, 1705060) (Biorad, 1705062).

For activation experiments: 2.5×10^4^ BV2 cells, 1.5×10^5^ NSCG, and 1×10^5^ NFpp10 cells were plated in 6-well plates (VWR, 734-1599) in their own media. After 48 hours, media was changed to 1% FBS DMEM for overnight incubation. Next day, conditioned media from BV2, NSCG and/or NFpp10 cells produced as described before during 72 hours in 1% FBS DMEM, was added on the BV2, NSCG and/or NFpp10 cells for 10 min, 30 min, 1 hour, 2 hours, or 6 hours.

For autophagy experiments: 2.5×10^4^ BV2 cells were plated in 6-well plates (VWR, 734-1599) in their own media. After 72 hours, BV2 cells were treated for 6 hours with either 2.5µM PLX3397 (MedChemExpress, HY-16749), 50nM Bafilomycin (Cayman Chemical, 11038) or both in combination.

For nuclear/cytoplasmic protein extraction: 1.5×10^5^ NSCG, and 1.5×10^5^ NFpp10 cells were plated in 6-well plates (VWR, 734-1599) in their own media. 2.5×10^4^ BV2 cells were plated in 6-well transwells (VWR, 734-2720) in their own media. After 48 hours, all the cells were changed to 10% FBS DMEM media and 50µM LPA 18:1 (Sanbio, 10010093-5) and/or 50µM LPA 16:0 (Sanbio, 10010093-5) were incubated overnight in 10% FBS DMEM at 4°C in a steering wheel. Next day, NSCG and NFpp10 cells were co-culture with BV2 cells or treated with the respective LPAs for 5 min, 10 min or 20 min. Protein extraction was performed using the Nuclear and Cytoplasmic Extraction Reagents (Thermofisher, 78833). For the nuclear fraction 2-5ug of protein were loaded among all conditions. For cytoplasmic fraction 50ug of protein were loaded among all conditions.

For RhoA pulldown experiments: 1.5×10^5^ NSCG, and 1×10^5^ NFpp10 cells were plated in 6-well plates (VWR, 734-1599) in their own media. 2.5×10^4^ BV2 cells were plated in 6-well transwells (VWR, 734-2720) in their own media. After 48 hours, all the cells were changed to 10% FBS DMEM media and 50µM LPA 18:1 (Sanbio, 10010093-5) and/or 50µM LPA 16:0 (Sanbio, 10010093-5) were incubated overnight in 10% FBS DMEM at 4°C in a steering wheel. Next day, NSCG and NFpp10 cells were co-culture with BV2 cells or treated with the respective LPAs for 2 min, 5 min or 10 min. Protein extraction was performed using the RhoA Pulldown activation Assay Kit (Cytoskeleton, BK036). For the pulldown 300-500ug of protein were loaded among all conditions. For the total fraction 20ug of protein were loaded among all conditions.

### Bulk lipidomics analysis

<u>For *in vitro* cells and media:</u> 2.5×10^4^ BV2 cells were plated in 6-well plates (VWR, 734-1599) in single and co-culture conditions with 1.5×10^5^ NSCG and 1×10^5^ NFpp10 cells using 6-well transwells (VWR, 734-2720) in 10% FBS DMEM. 2.5µM PLX3397 (MedChemExpress, HY-16749) was added when necessary. After 72 hours, cell and media samples were recollected. Cell samples were washed with ice-cold PBS and fresh-frozen in dry ice. Cell media samples were recollected, centrifuged to remove any cell debris, and fresh-frozen in dry ice.

<u>For *in vivo* tumor tissue:</u> fresh-frozen NSCG and NFpp10 tumor tissues were cold-PBS perfused. The brains were carefully extracted, cut at the injection point, and embedded in 3% carboxymethylcellulose (Merck, C4888). Tissues were fresh-frozen in pre-chilled isopentane (Acros organics, 126470010) using liquid nitrogen and stored at −80°C until further processing. For laser capture microdissection, 10µm sections were obtained using the Cryostar NX70 cryostat (Thermofisher), and mounted on FrameSlides with a 1.4µm PET membrane (Leica Microsystems, 11505151). After mounting, slides were dried in a vacuum desiccator for ∼10 minutes, frozen on dry ice, and finally stored at −80 °C. Slides were stained in an aqueous hematoxylin solution for 3 minutes. After washing with tap water, slides were dried again under vacuum. Selected tumor rim regions were extracted using a Leica Laser Microdissection (LMD) 6000B microscope equipped with a diode-pumped, solid-state laser at the wavelength of 355 nm, an average pulse energy of 70µJ, and a repetition rate of 80Hz. Regions of interest were selected to obtain a total of 5 mm² across five consecutive sections, using brightfield contrast, and captured in 0.2ml PCR tube caps (Greiner). The material was stored at −80°C until further processing.

<u>-Lipid extraction:</u> tissue samples were homogenized in microtubes with 200 µL of methanol for 30 min using a sonication bath. The lysate was transferred to glass Pyrex tubes and was mixed with 500 μl methanol, 100μl HCl(1M), 700μl water, 900μl CHCl3, 200μg/ml of the antioxidant 2,6-di-tert-butyl-4-methylphenol (BHT, Sigma Aldrich) and 3μl of UltimateSPLASH™ ONE internal standard mix (Avanti Polar Lipids, 330820) and 3μl of SphingoSPLASH™ I internal standard mix (Avanti Polar Lipids, 330734).

For cell pellet samples, the cell pellet was resuspended in 700μl water, transferred to glass Pyrex tubes and was mixed with 700μl methanol, 100μl HCl(1M), 900μl CHCl3 and antioxidant and internal standards as described above.

For media samples, 250μl medium was mixed with 450μl water, transferred to glass Pyrex tubes and was mixed with 700μl methanol, 100μl HCl(1M), 900μl CHCl3 and antioxidant and internal standards as described above. After vortexing and centrifugation, the lower organic fraction was collected and evaporated using a Savant Speedvac spd111v (Thermo Fisher Scientific) at room temperature and the remaining lipid pellet was stored at −20°C under argon. LPA was extracted as recently described^26^. Briefly, in glass Pyrex tubes, cells or medium were mixed with 1mL 0.1M HCl (in water), 1mL methanol, 3µL LPA 17:0 (10μg/mL stock, Avanti Polar Lipids, A85071). The mixture was vortexed for 2min, followed by the addition of 1 mL chloroform, another 2 min of vortexing and centrifugation at 5000g for 10min. The lower organic fraction was collected and evaporated using a Savant Speedvac, was reconstituted in 60μL methanol and transferred to an LC vial.

<u>-Mass spectrometry:</u> right before mass spectrometry analysis, lipid pellets were reconstituted in 100% ethanol. Lipid species were analyzed by liquid chromatography electrospray ionization tandem mass spectrometry (LC-ESI/MS/MS) on a Nexera X2 UHPLC system (Shimadzu) coupled with hybrid triple quadrupole/linear ion trap mass spectrometer (6500+ QTRAP system; AB SCIEX). Chromatographic separation was performed on a XBridge amide column (150mm × 4.6mm, 3.5μm; Waters) maintained at 35°C using mobile phase A [1mM ammonium acetate in water-acetonitrile 5:95 (v/v)] and mobile phase B [1mM ammonium acetate in water-acetonitrile 50:50 (v/v)] in the following gradient: (0-6 min: 0% B -> 6% B; 6-10 min: 6% B -> 25% B; 10-11 min: 25% B -> 98% B; 11-13 min: 98% B -> 100% B; 13-19 min: 100% B; 19-24 min: 0% B) at a flow rate of 0.7mL/min which was increased to 1.5mL/min from 13 minutes onwards. SM, CE, CER, DCER, HCER, LCER were measured in positive ion mode with a product ion of 184.1, 369.4, 264.4, 266.4, 264.4 and 264.4 respectively. TAG, DAG and MAG were measured in positive ion mode with a neutral loss for one of the fatty acyl moieties. PC, LPC, PE, LPE, PG, PI and PS were measured in negative ion mode by fatty acyl fragment ions. Lipid quantification was performed by scheduled multiple reactions monitoring (MRM), the transitions being based on the neutral losses or the typical product ions as described above. The instrument parameters were as follows: Curtain Gas = 35 psi; Collision Gas = 8 a.u. (medium); IonSpray Voltage = 5500 V and −4,500 V; Temperature = 550°C; Ion Source Gas 1 = 50 psi; Ion Source Gas 2 = 60 psi; Declustering Potential = 60 V and −80 V; Entrance Potential = 10 V and −10 V; Collision Cell Exit Potential = 15 V and −15 V. The LPA measurement method was adopted as recently described^69^. Measurements were performed on a Shimadzu Nexera X2 and Sciex QTRAP 6500+. Chromatographic separation was performed on a Nucleodur C8 Gravity column (125mm × 2.0mm, 5μm; Machery-Nagel) maintained at 30°C using mobile phase A [methanol:water 75:25 (v/v) with 0.5% formic acid and 5 mM ammonium formate] and mobile phase B 80% [methanol:water 99:1 (v/v) with 0.5% formic acid and 5 mM ammonium formate] + 20% Chloroform, in the following gradient: (0-1 min: 0% B; 1-11 min: 0% B -> 100% B; 11-15 min: 100% B; 15-16 min: 100% B -> 0% B; 16-21 min: 0% B; at a flow rate of 0.5 mL/min. LPA was measured in negative ion mode with MRM transitions to fragment 153.0, with instrument parameters as follows: Curtain Gas = 30 psi; Collision Gas = 8 a.u. (medium); IonSpray Voltage = −4,500 V; Temperature = 300°C; Ion Source Gas 1 = 60 psi; Ion Source Gas 2 = 45 psi; Declustering Potential = −90 V; Entrance Potential = −10 V; Collision Cell Exit Potential −15 V and Collision Energy −26 V. LPA retention times were distinct from LPC and identifications were curated using the equivalent carbon number (ECN) retention time model.

The following fatty acyl moieties were taken into account for the lipidomic analysis: 14:0, 14:1, 16:0, 16:1, 16:2, 18:0, 18:1, 18:2, 18:3, 20:0, 20:1, 20:2, 20:3, 20:4, 20:5, 22:0, 22:1, 22:2, 22:4, 22:5 and 22:6 except for TGs which considered: 16:0, 16:1, 18:0, 18:1, 18:2, 18:3, 20:3, 20:4, 20:5, 22:2, 22:3, 22:4, 22:5, 22:6.

<u>-Data Analysis:</u> peak integration was performed with the MultiQuantTM software version 3.0.3. Lipid species signals were corrected for isotopic contributions (calculated with Python Molmass 2019.1.1) and were quantified based on internal standard signals and adheres to the guidelines of the Lipidomics Standards Initiative (LSI) (level 2 type quantification as defined by the LSI).

### Quantitative Mass Spectrometry Imaging (qMSI)

NSCG and NFpp10 tumor tissues were cold-PBS perfused. The brains were carefully extracted, cut at the injection point, and embedded in 3% Carboxymethylcellulose (Merck, C4888). Tissues were fresh-frozen in pre-chilled isopentane (Acros organics, 126470010) using liquid nitrogen and stored at −80 °C until further processing. Ten µm sections were obtained using the Cryostar NX70 cryostat (Thermofisher), and mounted on IntelliSlides (Bruker Daltonik GmbH). Slides were then dried using a vacuum desiccator for 30 min and scanned using an FS120 slide scanner (Braun, Germany).

<u>-Application of internal standards and matrix:</u> an in-house internal lipid standard mixture was prepared with 0.100mg/ml PE 15:0/18:1 [D7] (Avanti Polar Lipids, 791638), 0.161mg/ml PC [D5] 17:0/16:1 (Avanti Polar Lipids, 855682), 0.161mg/ml PC [D9] 16:0/16:0 (Avanti Polar Lipids, 860352), 0.031mg/ml SM d18:1;O2/18:1 [D9] (Avanti Polar Lipids, 791649), 0.033mg/ml Cer 18:1;O2/12:0 (Cayman Chemical, 22530), 0.0030mg/ml LPC [D9] 16:0 (Echelon Biosciences, L-1516D), 0.0030mg/ml LPC 18:1 [D7] (Avanti Polar Lipids, 791643), 0.0030mg/ml LPC [D9] 18:0 (Echelon Biosciences, L-1518D) and 0.00025mg/ml L-CAR [D9] 16:0 (Avanti Polar Lipids, 870323), and was diluted 8x in methanol for spraying performed using an HTX M5 sprayer (HTX Imaging, HTX Technologies) connected with a KD Legato 100 Infuse Single pump (KdScientific) according to a published protocol^70^. After spraying the internal standard mixture, a peristaltic pump (Azure P4.1S) was connected to the sprayer to apply the matrix. A solution of 5 mg/ml norharmane (Sigma-Aldrich) in chloroform:methanol (2:1 v/v) was prepared and sonicated for 15 minutes. The matrix was filtered before application through the 2µm MillexLG 13 mm PhilicPTFE NS filter. Ten layers were sprayed in the CC pattern, with a track spacing of 2mm, at a nozzle temperature of 30°C and a plate temperature of 25°C, a flow rate of 120µl/min, a velocity of 1350mm/min, a nozzle height of 40mm, and 10psi nitrogen pressure.

<u>-Mass Spectrometry Imaging parameters:</u> MSI was performed using a timsTOF fleX MALDI-2 mass spectrometer (Bruker Daltonik) equipped with a smartbeam 3D 10=kHz laser. Data were acquired in positive-ion mode with a 16×16 µm scan range, yielding a 20×20 µm pixel size. The MALDI laser was operated at 65% laser power in single smart-beam mode, with beam scanning enabled, at 250 shots per pixel at 10 kHz laser trigger frequency. The *m/z* range was 300-1800. The MALDI plate offset was set to 50 V, and the deflection delta to 70 V. The funnel 1 and 2 RF voltages were 300 and 350 Vpp, respectively, and the multipole RF was 400 Vpp. The energy difference between multipole and quadrupole was 5 eV, with the lowest mass on the quadrupole set to *m/z* 300. The energy difference between the quadrupole and the collision cell (collision energy) was 10 eV, and the collision cell RF was 1800Vpp. The focus pre-TOF settings, including transfer time to the TOF stage and pre-pulse storage, were set to 80µs and 12µs.

<u>-Data Analysis:</u> all collected data was initially imported into SCiLS lab MVS (v2026b Pro) using standard import settings, converted to .imzML files, and subsequently processed in the LipidQMap software platform (v0.2.0-beta 7) into quantitative mass spectrometry (QMSI) images^71^. Raw data was imported with a mass accuracy of 10ppm and a bin size of 5mDa, and missing pixels were interpolated using the median value of the surrounding 3 x 3-pixel neighborhood. Sodium isobaric overlaps were corrected as previously described^72^, and type II isobaric corrections were applied to account for double-bond–related isobaric overlap. Data was recalibrated using the PC 34:1 [M+K]^+^ adduct as the reference, with a peak assignment tolerance of 30 ppm and a minimum intensity threshold of 10000 required for recalibration. Lipid adducts were annotated using an in-house Python-generated database of lipids (available within LipidQMap software). The raw, corrected, and QMSI images were winsorized at the 98th percentile. For LPC 16:0, 18:0, and 18:1, the [M+H]^+^, [M+K]^+^, and sum of these two adducts were exported as .png files. In the next step, the standard normalized data (as summed adducts) were reimported into the SCiLS platform through the API. Bisecting k-means clustering was performed using the correlation distance as the metric.

### MILAN multiplex immunohistochemistry of human brain sections

<u>-Tissue staining:</u> multiplex immunohistochemistry was performed in 50 tissue microarrays (TMAs) coming from 18 different glioblastoma patients (collected at the UZ/KU Leuven biobank under protocol S-62248) according to the previously published method^73,74^. Three µm formalin-fixed paraffin-embedded (FFPE) tissue sections were dewaxed, and antigen retrieval was performed using PT link (Agilent) using 10mM EDTA in Tris-buffer pH 8. Immunofluorescent staining was performed using the Bond RX Fully Automated Research Stainer (Leica biosystems) and the primary antibodies listed in **Extended Data Table 1**. Tissue sections were incubated during 4 hours with the primary antibodies, washed and visualized with secondary antibodies (30 minutes, **Extended Data Table 2**). A coverslip was placed on the slides with medium containing DAPI and were scanned in the Zeiss Axio Scan Z.1 (Zeiss) with 10X magnification. After imaging, coverslips were manually removed after 30 minutes soaking in washing buffer. Consecutive washing steps were thereafter performed using TBS. Stripping of the antibodies was performed in a buffer containing 1% SDS/β-mercaptoethanol during 30 minutes at 56°C. After the stripping procedure, slides were washed in washing buffer for 45 minutes with frequent buffer changes. This cycle repeated across multiple marker combinations.

<u>-Image processing:</u> raw scans were transformed to grayscale 16-bit TIFF images using developer’s software (ZEN). Image registration was performed using the imreg package implemented in Python. Autofluorescence (AF) subtraction was performed by subtracting the pre-stained image of the corresponding tissue section from the measured signal (MS). Intensities for both images were normalized using quantile normalization. Cellular segmentation was done by applying STARDIST^75^ on the images from DAPI channel. This segmentation technique accurately delineates contours for each cell present in the tissue. For each cellular object, morphological (nuclear size) and functional (marker intensity) features were extracted, following removal of the objects that overlapped with strong auto fluorescent regions.

<u>-Data analysis:</u> phenotypic identification was implemented as previously described^74^. Briefly, three different clustering methods were applied (PhenoGraph, FlowSom, and KMeans), from which only those cells that agreed over at least 2 algorithms using a consensus-based approach were retained. Selected clusters were then manually annotated and assigned to the right cell type. First level clustering was used to define the main cell types (tumor cells, macrophages, microglia, T cells and blood vessels), following which in each category subclusters were defined as outlined in the paper.

### Single-cell RNA sequencing and analysis

NSCG and NFpp10 tumor rim and core samples were dissected under a fluorescence stereo microscope and digested using collagenases and DNase, and single-cell suspensions were processed with 10X Genomics Chromium (targeting 6,000 cells/library). Libraries were sequenced on Illumina HiSeq4000 and aligned to the mouse genome (mm10) using CellRanger v7.0.1. Data was processed in Seurat5 with lenient filtering to retain low-transcription cells (e.g., basophils). Data was normalized, centered, and scaled, and the CHOIR pipeline^76^ was used to perform PCA and UMAP dimensionality reduction and generate initial Leiden clustering. Harmony was used to correct for batch effects. Cluster resolution was further refined manually, and cluster annotations were based on canonical marker genes previously defined in the literature. Doublet clusters (defined based on dual expression of marker genes) were excluded from downstream analysis. Objects were converted to the *anndata* format for enrichment analysis via the decoupler (v2.1.4) package^77^ and scored via the ULM method using the GO Biological Processes, Reactome, and Hallmark pathway databases.

### Bulk RNA sequencing and analysis

1.5×10^5^ NSCG, and 1×10^5^ NFpp10 cells were plated in 6-well plates (VWR, 734-1599) in their own media. After 48 hours, tumor cells were treated with 50µM LPA 16:0 (Sanbio, 10010093-5), and/or 50µM LPA18:1 (Sanbio, 10010093-5) for 20 hours. As above-mentioned, LPAs were previously incubated overnight in 10% FBS DMEM at 4°C in a steering wheel. After 20 hours treatment, cell pellets were recollected and sent to Novogene for sequencing. Reads were aligned to the mouse genome (mm39) with Hisat2 (v2.0.5) and quantified with featureCounts (v1.5.0-p3). Differential expression analysis was performed using PyDESeq2 (v0.5.4), with differential genes defined as having Benjamini-Hochberg adjusted p-values < 0.01. Enrichment analysis was performed via the decoupler (v2.1.4) package and scored via the ULM method using the GO Biological Processes, Reactome, and Hallmark pathway databases.

### TCGA

The TCGA-GBM Pan-Cancer Atlas mRNA sequencing data were downloaded from cBioPortal on January 29, 2026 and visualized using GraphPad Prism.

### Spatial transcriptomics

OCT-embedded GBM tumor brains were cryosectioned on a CryoStar NX70 cryostat (Thermo Fisher Scientific) at 8µm thickness (chamber temperature −12=°C to −14=°C) before mounting on a MERSCOPE slide (Vizgen). Prepared sections were refrozen at −20=°C for 30=min, brought to room temperature for 7=min, and fixed in 4% paraformaldehyde (PFA) for 15=min at room temperature. Sections were washed three times with PBS, air-dried for 30=min, and permeabilized in 70% ethanol at 4=°C until processing.

Multiplexed Error-Robust Fluorescence In Situ Hybridization (MERFISH) was performed on the Vizgen MERSCOPE platform following the manufacturer’s protocol (550 mouse brain gene panel/codebook: CP1622, MERSCOPE software version: 234b.241217.1593). Briefly, the sample was washed with the conditioning buffer and incubated with the pre-anchoring buffer for 2h at 37°C, followed by a formamide buffer wash for 30min at 37°C. The sample was then incubated overnight at 37°C with the anchoring buffer. The section was washed with formamide buffer at 47°C, and then a gel was cast on top using the gel embedding solution (Vizgen with 10% APS and TEMED) covered with a Sigmacote-coated glass coverslip for 90min at room temperature. The glass coverslip was removed and the tissue was cleared in clearing premix with proteinase K for 2h at 47°C. The auto-fluorescence was quenched for 3h at room temperature in MERSCOPE Photobleacher in clearing solution without proteinase K. The sample was then incubated for 48h at 37°C with the MERSCOPE Gene Panel Mix (CP1622). Before imaging, the sample was washed with formamide wash buffer and sample prep wash buffer, then stained with DAPI and PolyT staining reagents for 15min at room temperature, washed again with formamide buffer and finally with sample prep wash buffer. The prepared sample was then immediately imaged in the Vizgen MERSCOPE instrument following the instrument user guide (91600001 RevK). All buffers and reagents were from the Vizgen kit unless otherwise indicated. Primary processing (image registration, barcode decoding, transcript calling, and cell segmentation) was performed using the MERSCOPE software pipeline, generating a cell-by-gene count matrix and spatial coordinates for detected transcripts. For downstream visualization, the cell-by-gene matrix and associated metadata were converted to a SCope-compatible .loom file and loaded into SCope [Davie et al., Cell (2018)]. Spatial expression maps for *Egfr*, *Tmem119*, and *Enpp2* were exported for figure generation.

### Statistical analysis

Statistical tests (t-test, Log-rank, Mann-Whitney, Wilcoxon) were performed using Prism and R. Significance was defined as p<0.05 with multiple comparison corrections as needed. Further details are provided in the Supplemental Material.

## Funding

-Fonds Wetenschappelijk Onderzoek Vlaanderen (FWO): project numbers 11F1423N (C.P-M), G0A1122N (G.B), S001623N SBO-LIPOMACS (J.V. S.), (S.-M.F).

-Kom op tegen Kanker (KOTK): grant number 13897 (C.P-M).

-The European Union’s Horizon Europe research and innovation programme under the Marie Skłodowska-Curie Doctoral Networks: grant agreements 766069 (GLIOTRAIN) (L.W), 101073386 (GLIORESOLVE) (K.S.G).

-Stichting Tegen Kanker (STK): grant numbers 2019-093 (M.G), 2018-086 FAF-F/2018/1303 (G.B), (S.-M.F.).

-Dieter de Cauderlier Fonds voor Hersentumoren (J.V.S).

-KU Leuven Opening the Future (J.V.S).

-Leuven Future Fund LISCO-Biomed (J.V.S).

-Leuven Cancer Institute (LKI) Focus Group Grant (J.V.S).

-Wereld Kanker Onderzoek Fonds (WKOF) (S.-M.F.).

-KU Leuven Methusalem (METH/26/009) (S.-M.F.)

## Acknowledgments

We kindly thank Kevin Feyen, Martine Nijs, Polina Zahdai, Lydia Boyle-García, Kathryn Jacobs, Dorothy John Robbert, Sophie Guelfi and Kan Lu for their help and technical assistance. We gratefully acknowledge the VIB Single Cell Core, and the VIB Nucleomics Core in Leuven.

## Author contributions

C.P-M, M.G and G.B designed most experiments. C.P-M, M.G, B.R, and L.W performed all experiments unless specified. J.I, J.D, N.R, and X.S performed and analyzed the bulk lipidomics and qMSI. K.A.N, and N.D.L performed and analyzed the MILAN multiplex immunohistochemistry. V. K analyzed the spatial transcriptomics data. K.S.G performed the bulkRNAseq and scRNAseq bioinformatic analysis. J.V.S, F.D.S, and S.-M.F provided critical input. J.V.S offered feedback on methodology, data curation, and assisted with funding acquisition. G.B and C.P-M conceptually planned the study. G.B supervised the study. C.P-M wrote the initial draft of the manuscript. G.B and C.P-M edited and amended the manuscript.

## Data availability

All data needed to evaluate the conclusions in the paper are present in the paper and/or the Supplementary Materials. The newly generated murine scRNA-seq and bulkRNA-seq data (fastq files and processed data) have been deposited in GEO. Spatial transcriptomics data is not currently available since it is part of another manuscript.

## Supplementary material

**Figure S1.**
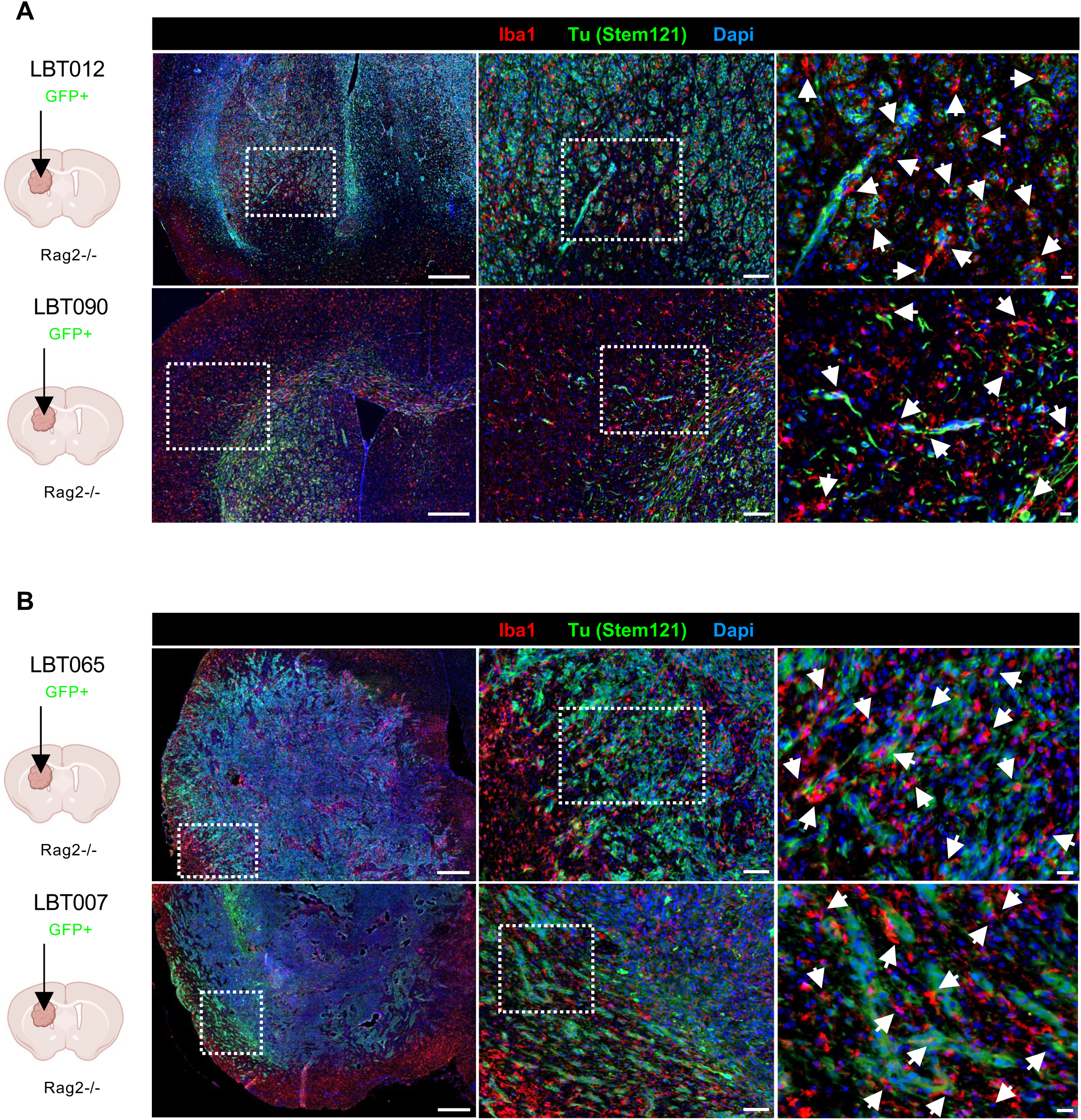
Spatial association of MGs and GBM patient-derived cell lines at the tumor rim. (A) Immunofluorescence images of LBT012 LamB (upper panel) and LBT090 Lami (lower panel) patient-derived GBM cells (EGFR amplified/muta^+^, Stem121^+^, green) associated with microglial cells (Iba1^+^, red) at the invasive rim of tumor-bearing mice. Scales (from left to right) 500µm, 100µm, and 20µm. (B) Immunofluorescence images of LBT065 LamB (upper panel) and LBT007 Lami (lower panel) patient-derived GBM cells (EGFR wildtype, Stem121^+^, green) associated with microglial cells (Iba1^+^, red) at the invasive rim of tumor-bearing mice. Scales (from left to right) 500µm, 100µm, and 20µm.

**Figure S2.**
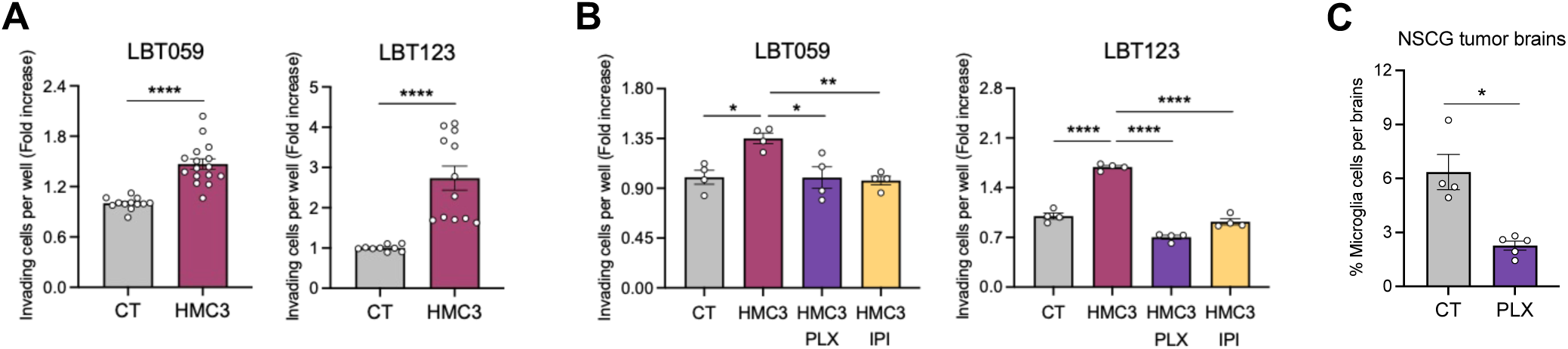
Human GBM invasion relies on the MG activation state. (A) Invasion assays of LBT059 Lami (left) and LBT123 LamB (right) cells (EGFR wildtype) in the presence or absence of human HMC3 microglial cells. Fold increase. LBT059 Lami: n=7, Mann-Whitney test (T-test non-parametric); **** p-value<0.0001; LBT123 LamB: n=5, Mann-Whitney test (T-test non-parametric), Mean with *±* SEM, ****p-value <0.0001. (B) Invasion assays of LBT059 Lami (left) and LBT123 LamB (right) cells (EGFR wildtype) in the presence or absence of human HMC3 microglial cells, PLX3397, and/or IPI-145. Fold increase. LBT059 Lami: n=4, One-way ANOVA, * p-value<0.0126, ** p-value=0.0072; LBT123 LamB: n=4, One-way ANOVA, Mean with *±* SEM, **** p-value<0.0001. (C) Microglia quantification in NSCG control- and PLX5622-treated tumor-bearing mice. n=4 control brains. n=5 PLX-treated brains. Mann-Whitney test (T-test non-parametric), * p-value=0.0159. All data represent mean ± SEM.

**Figure S3.**
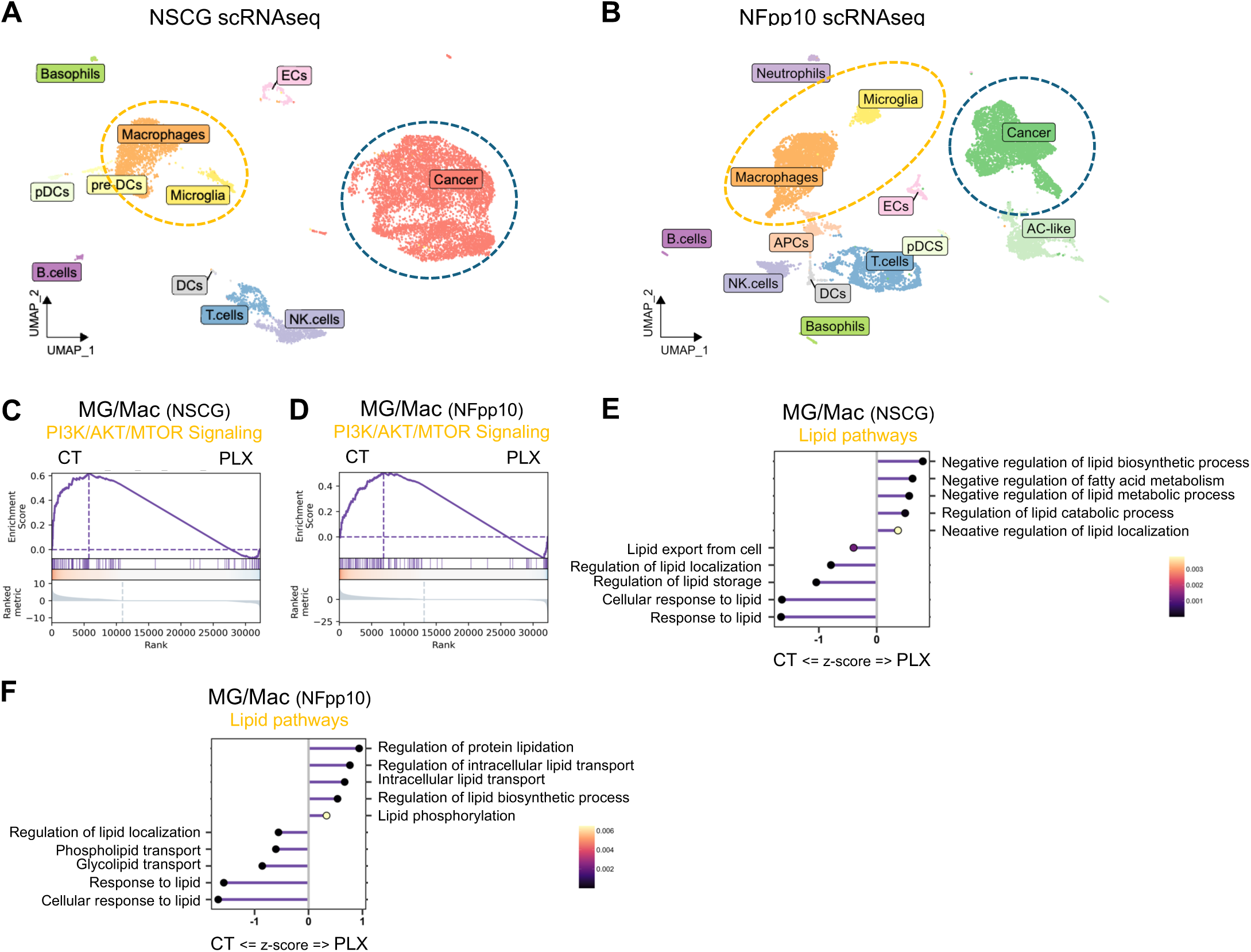
GBM-associated MG undergo lipid metabolic reprograming. (A) UMAP plot of the different cell clusters in NSCG-tumor brains. (B) UMAP plot of the different cell clusters in NFpp10-tumor brains. (C) Leading edge plot of the PI3K/AKT/MTOR signaling hallmark in microglia/macrophages of NSCG tumor rim control and PLX-treated samples. p-adj=8.17E-45. NES= 1.686654. (D) Leading edge plot of the PI3K/AKT/MTOR signaling hallmark in microglia/macrophages of NFpp10 tumor rim control and PLX-treated samples. p-adj=6.97E-49. NES= 1.205815. (E) Lollipop chart of the most significant lipid pathways in microglia/macrophages of NSCG tumor rim control and PLX-treated samples. (F) Lollipop chart of the most significant lipid pathways in microglia/macrophages of NFpp10 tumor rim control and PLX-treated samples.

**Figure S4.**
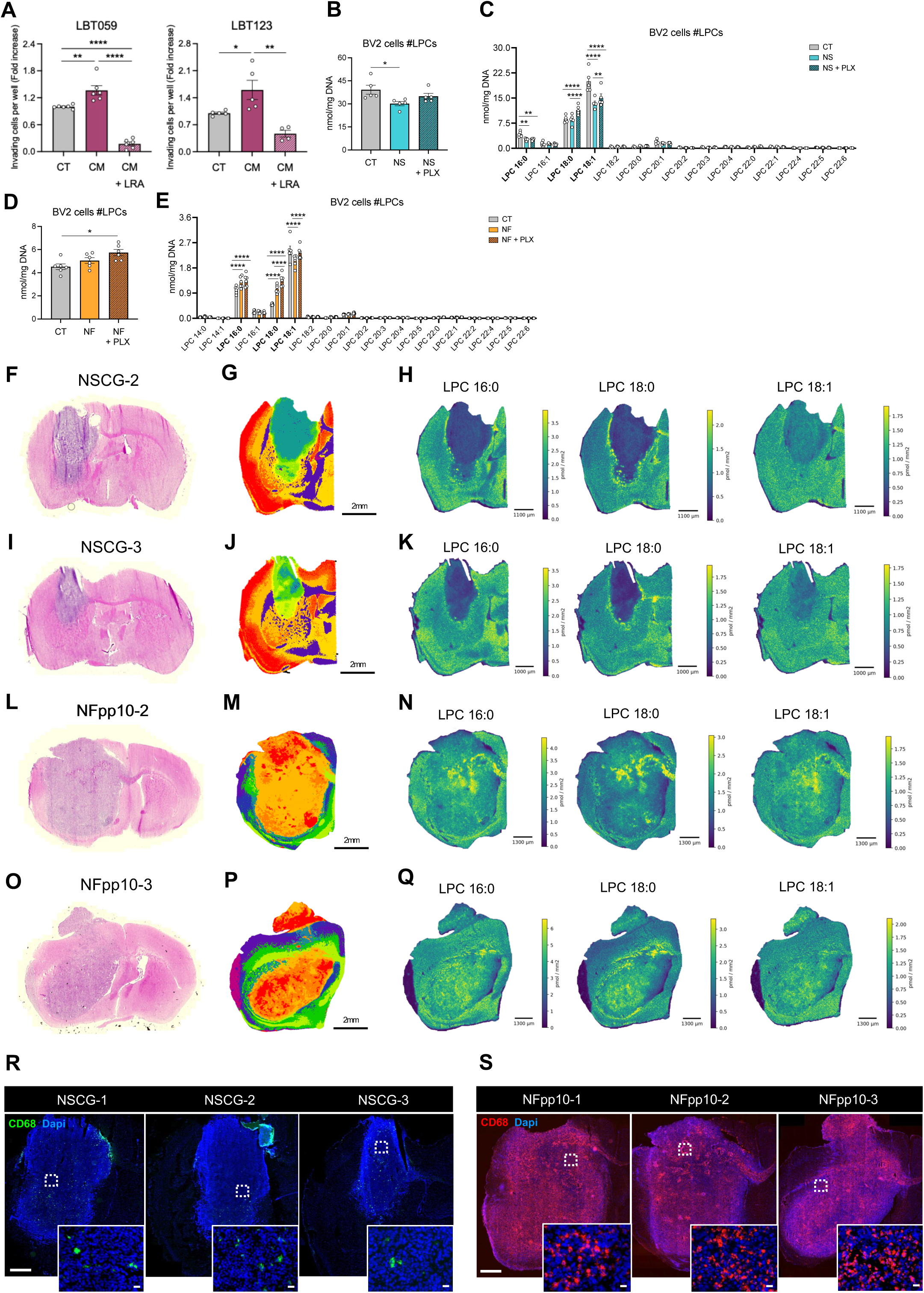
MG-secreted lipids drive GBM invasion. (A) Invasion assays of LBT059 Lami (left) and LBT123 LamB (right) cells (EGFR wildtype) in the presence or absence of human HMC3 conditioned-media (CM) with or without lipid removal agent (LRA). Fold increase. LBT059 Lami: n=3, One-way ANOVA, ** p-value=0.0036, **** p-value<0.0001; LBT123: n=3, One-way ANOVA, * p-value=0.0491, ** p-value=0.0014. (B) Quantification of the sum of LPC species in BV2 cells in the absence or presence of NSCG co-culture and/or PLX3397. n=5. One-way ANOVA, * p-value=0.0318. (C) Quantification of the distinct LPC species in BV2 cells in the absence or presence of NSCG co-culture and/or PLX3397. n=5. Two-way ANOVA, ** p-value<0.0641, ****p-value<0.0001. (D) Quantification of the sum of LPC species in BV2 cells in the absence or presence of NFpp10 co-culture and/or PLX3397. n=6. One-way ANOVA, ** p-value=0.0022. (E) Quantification of the distinct LPC species in BV2 cells in the absence or presence of NFpp10 co-culture and/or PLX3397. n=6. Two-way ANOVA, **** p-value<0.0001. (F) Hematoxylin & Eosin (H&E) staining of an NSCG tumor brain (NSCG-2). (G) K-means lipid clustering of LPC, PC, PC P-/PC O-, SM, CAR in a NSCG tumor brain sample (NSCG-2) showing different lipid profiles in distinct areas. Signals normalized to their internal standard. Each color represent one lipid cluster. Scale 2mm. (H) Quantitative Mass Spectrometry Imaging (qMSI) of LPC 16:0, LPC 18:0, and LPC 18:1 in an NSCG tumor brain (NSCG-2). Scale 1100µm. (I) Hematoxylin & Eosin (H&E) staining of an NSCG tumor brain (NSCG-3). (J) K-means lipid clustering of LPC, PC, PC P-/PC O-, SM, CAR in a NSCG tumor brain sample (NSCG-3) showing different lipid profiles in distinct areas. Signals normalized to their internal standard. Each color represent one lipid cluster. Scale 2mm. (K) Quantitative Mass Spectrometry Imaging (qMSI) of LPC 16:0, LPC 18:0, and LPC 18:1 in an NSCG tumor brain (NSCG-3). Scale 1000µm. (L) Hematoxylin & Eosin (H&E) staining of an NFpp10 tumor brain (NFpp10-2). (M) K-means lipid clustering of LPC, PC, PC P-/PC O-, SM, CAR in a NFpp10 tumor brain sample (NFpp10-2) showing different lipid profiles in distinct areas. Signals normalized to their internal standard. Each color represent one lipid cluster. Scale 2mm. (N) Quantitative Mass Spectrometry Imaging (qMSI) of LPC 16:0, LPC 18:0, and LPC 18:1 in an NFpp10 tumor brain (NFpp10-2). Scale 1300µm. (O) Hematoxylin & Eosin (H&E) staining of an NFpp10 tumor brain (NFpp10-3). (P) K-means lipid clustering of LPC, PC, PC P-/PC O-, SM, CAR in a NFpp10 tumor brain sample (NFpp10-3) showing different lipid profiles in distinct areas. Signals normalized to their internal standard. Each color represent one lipid cluster. Scale 2mm. (Q) Quantitative Mass Spectrometry Imaging (qMSI) of LPC 16:0, LPC 18:0, and LPC 18:1 in an NFpp10 tumor brain (NFpp10-3). Scale 1300µm. (R) Immunofluorescence images of the NSCG tumor samples used in the quantitative Mass Spectrometry Imaging (qMSI) analysis focusing on the microglia/macrophages (CD68, green). Scale upper to lower: 1000µm, 20µm. (S) Immunofluorescence images of the NFpp10 tumor samples used in the quantitative Mass Spectrometry Imaging (qMSI) analysis focusing on the microglia/macrophages (CD68, red). Scale upper to lower: 1000µm, 20µm. All data represent mean ± SEM.

**Figure S5.**
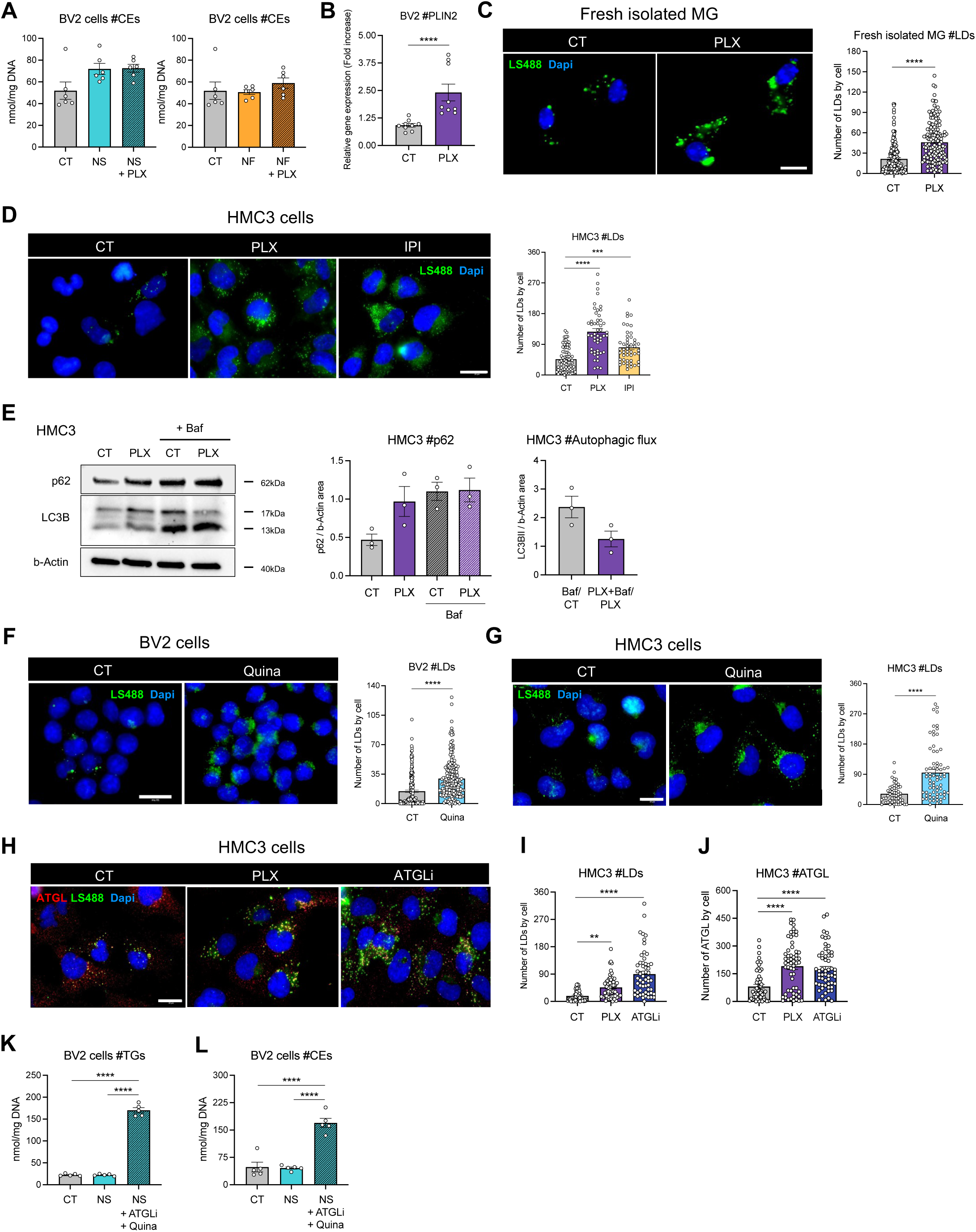
MG-CSF1R and PI3Kγ/δ inhibition blocks lipophagy/lipolysis, promotes LD accumulation and impairs GBM invasion. (A) Quantification of the sum of CE species in BV2 cells in the absence or presence of NSCG co-culture (left) or NFpp10 co-culture (right) with or without PLX3397. BV2 (NSCG co-culture): n=6, One-way ANOVA, non-significant. BV2 (NFpp10 co-culture): n=6, One-way ANOVA, non-significant. (B) Gene expression of PLIN2 (Perilipin-2) in BV2 cells untreated or PLX3397-treated. Relative gene expression to S18. Fold increase. n=3. Mann-Whitney T-test (non-parametric), **** p-value<0.0001. (C) Immunofluorescence images and corresponding quantification of freshly isolated murine microglial cells (Dapi, blue) and LDs (LS488, green) upon PLX3397 treatment. Scale bar: 10µm. n=3. Mann-Whitney test, **** p-value<0.0001. (D) Immunofluorescence images and corresponding quantification of HMC3 cells (Dapi, blue) and LDs (LS488, green) upon PLX3397 and IPI-145 treatment. Scale bar: 20µm. n=3. One-way ANOVA, *** p-value<0.0004, **** p-value<0.0001. (E) Representative immunoblots and quantifications of p62, and LC3B levels in HMC3 cells untreated or treated with PLX3397 in the absence or presence of bafilomycin (Baf). n=3. One-way ANOVA, non-significant. (F) Immunofluorescence images and corresponding quantification of BV2 cells (Dapi, blue) and LDs (LS488, green) upon quinacrine (Quina) treatment. Scale bar: 20µm. n=5. Mann-Whitney T-test; **** p-value<0.0001. (G) Immunofluorescence images and corresponding quantification of HMC3 cells (Dapi, blue) and LDs (LS488, green) upon quinacrine (Quina) treatment. Scale bar: 20µm. n=4. Mann-Whitney T-test, **** p-value<0.0001. (H, I, J) Immunofluorescence images (H) and corresponding quantification of HMC3 cells (Dapi, blue), LDs (LS488, green) (I), and ATGL (ATGL, red) (J) upon PLX3397 and ATGL inhibitor (ATGLi) treatment. Scale bar: 20µm. HMC3 – LDs (I): n=3, One-way ANOVA, ** p-value=0.0063, **** p-value<0.0001; HMC3 – ATGL (J): n=3, One-way ANOVA, **** p-value<0.0001. (K) Quantification of the sum of TG species in BV2 cells in the absence or presence of NSCG co-culture with or without ATGLi and quinacrine. n=5, One-way ANOVA, **** p-value<0.0001. **l.** Quantification of the sum of CE species in BV2 cells in the absence or presence of NSCG co-culture with or without ATGLi and quinacrine. n=5, One-way ANOVA, **** p-value<0.0001. All data represent mean ± SEM.

**Figure S6.**
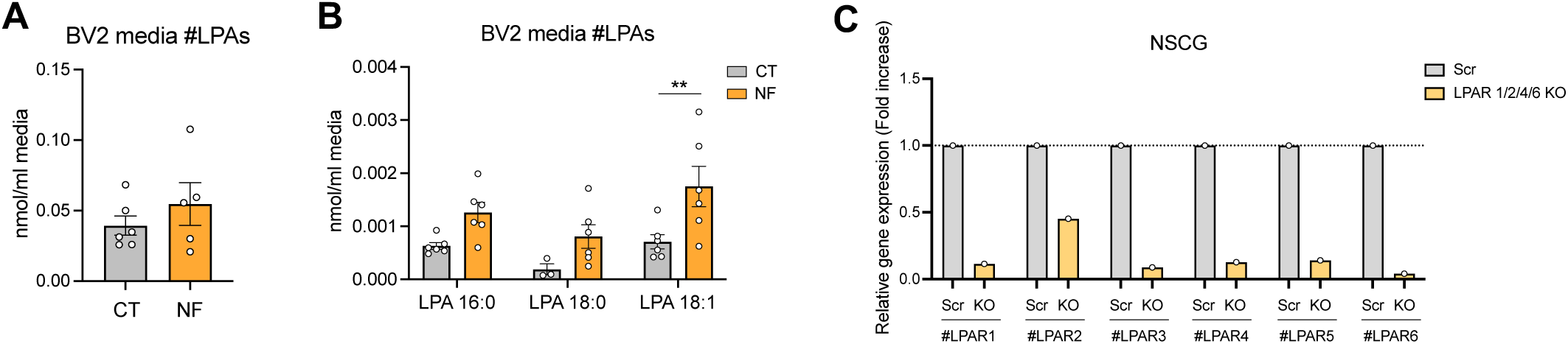
LPA-LPAR signaling promotes GBM invasion. (A) Quantification of the sum of LPA species in BV2 media in the absence or presence of NFpp10 co-culture. n=5. Mann-Whitney test, non-significant. (B) Quantification of the distinct LPA species in BV2 media in the absence or presence of NFpp10 co-culture. n=5. Two-way ANOVA, ** p-value=0.0065. (C) Gene expression of the LPAR1-6 of the NSCG Scr (control) and NSCG LPAR1/2/4/6 Knock-out cells injected *in vivo*. Relative gene expression to S18. Fold increase. n=1. All data represent mean ± SEM.

**Figure S7.**
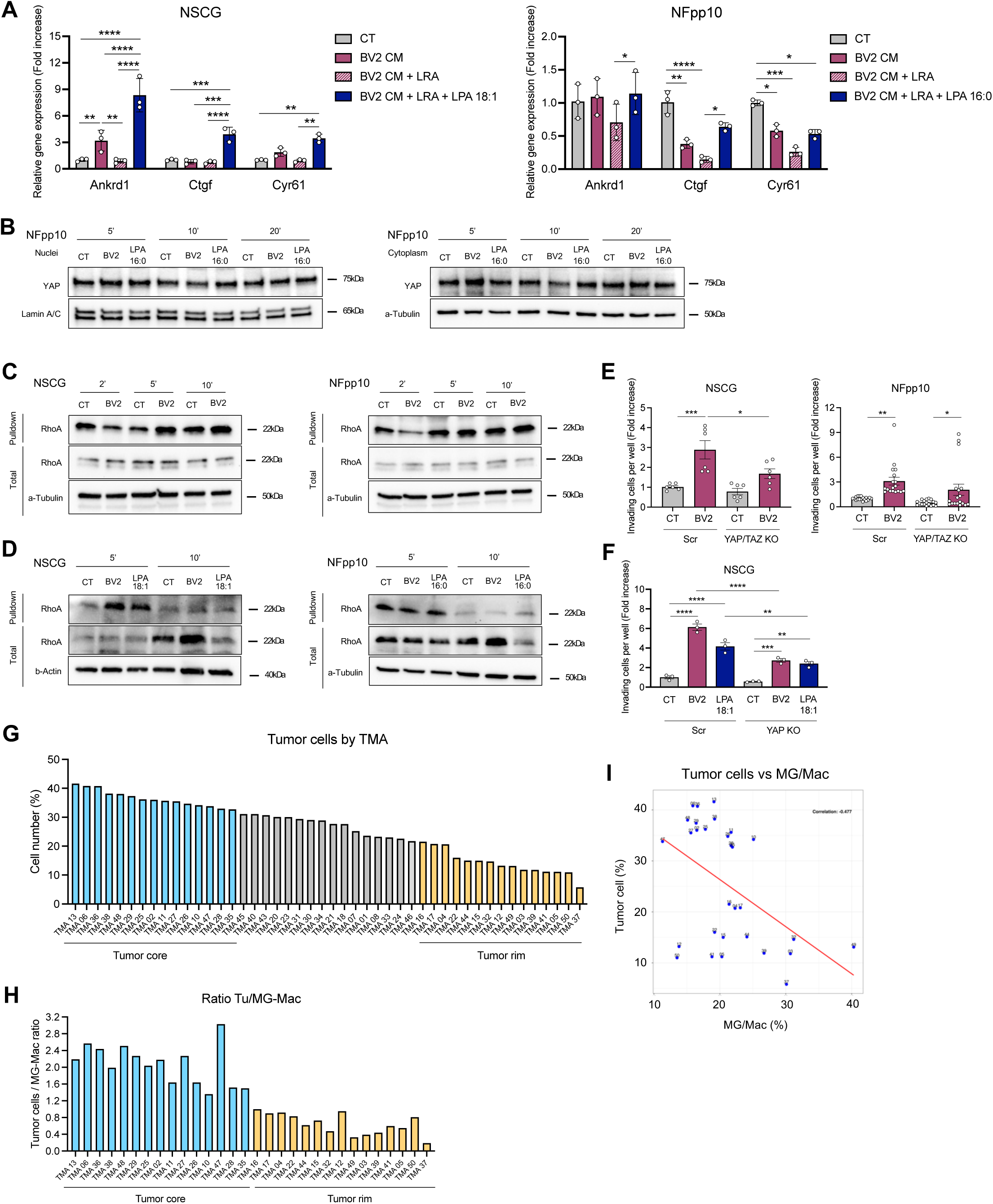
EGFRvIII^+^ GBM invasion relies on Yap/Taz activation. (A) Gene expression of Yap-target genes (*Ankrd1, Ctgf, Cyr61*) in NSCG (left) and NFpp10 (right) cells upon 24 hours of treatment with BV2 CM, BV2 CM LRA (lipid removal), BV2 CM LRA (lipid removal) LPA 18:1. Relative gene expression to S18. Fold increase. NSCG: n=3, 2way ANOVA, ** p-value<0.0056; *** p-value<0.0003, **** p-value<0.0001; NFpp10: n=3, 2way ANOVA, * p-value<0.0381, ** p-value=0.0012, *** p-value=0.0002, **** p-value<0.0001. (B) Representative immunoblots of Yap nuclear and cytoplasmic levels in NFpp10 cells upon 5 minutes, 10 minutes, 20 minutes of co-culture with BV2 cells and LPA 16:0 treatment. (C) Representative immunoblots of RhoA pulldown and total in NSCG (left) and NFpp10 (right) cells upon 2 minutes, 5 minutes, 10 minutes co-culture with BV2 cells. (D) Representative immunoblots of RhoA pulldown and total in NSCG (left) and NFpp10 (right) cells upon 5 minutes, 10 minutes co-culture with BV2 cells and LPA 18:1 or LPA 16:0 treatment respectively. (E) Invasion assays of NSCG (left) and NFpp10 (right) Scr (control) and YAP/TAZ knock-out cells in the presence or absence of BV2 cells. Fold increase. NSCG: n=3, One-way ANOVA, * p-value=0.0267, *** p-value=0.0001; NFpp10: n=6, One-way ANOVA, * p-value=0.0432, ** p-value=0.0034. (F) Invasion assays of NSCG Scr (control) and YAP knock-out cells in the presence or absence of BV2 cells and LPA 18:1 treatment. Fold increase. NSCG: n=3, One-way ANOVA, ** p-value<0.0022, *** p-value=0.0004, **** p-value<0.0001. (G) Tumor cell number across the 50 distinct TMAs. **h.** Ratio tumor cell/microglia-macrophages across the selected 15 tumor core and 15 tumor rim TMAs based on tumor cell density. **i.** Correlation tumor cells/microglia-macrophages across the selected 15 tumor core and 15 tumor rim TMAs. Each dot/number refers to a TMA sample. Correlation index: −0.477. All data represent mean ± SEM.

